# Zygotic genome activation is timed by the egg DNA-to-cytoplasm ratio across animals

**DOI:** 10.64898/2026.04.16.718233

**Authors:** Israel Campo-Bes, Federica Mantica, Jon Permanyer, Cristina Rodríguez-Marin, Kero Guynes, Tòt Senar-Serra, Gonzalo Quiroga-Artigas, Yan Liang, Allan M. Carrillo-Baltodano, Josefa Cruz, Rossella Annunziata, Sandra Chevalier, Marta Iglesias, Fumiaki Sugahara, Yi-Jyun Luo, Anna Schoenauer, Jorge Corbacho, Periklis Paganos, Filomena Caccavale, Rosa Maria Sepe, Luis P. Iñiguez, Mette Handberg-Thorsager, Christina Zakas, Ralf J. Sommer, Maria Ina Arnone, Alistair P. McGregor, Juan R. Martínez-Morales, Noriyuki Satoh, Hector Escriva, Stéphanie Bertrand, Arnau Sebé-Pedrós, Enrico D’Aniello, Juan Pascual-Anaya, Evelyn Houliston, Xavier Franch-Marro, David Martín, Ildiko M.L. Somorjai, Isabel Almudi, José M Martín-Durán, Yi-Hsien Su, Alejandro Burga, Salvatore D’Aniello, Manuel Irimia

## Abstract

During early animal embryogenesis, gene expression control shifts from maternally deposited products to newly transcribed RNA through zygotic genome activation (ZGA). Although essential and universal, ZGA remains poorly understood outside a few model species. Here, we generated a transcriptomic atlas of early embryogenesis across 61 animal species from 13 phyla and used a unified computational framework to infer ZGA timing systematically. We find extensive variation in ZGA onset across animals, but show that the ratio between genome size and egg cytoplasmic volume, a proxy for the nuclear-to-cytoplasmic ratio, robustly predicts when genome activation begins. We also reveal that zygotic genes also differ from maternal transcripts in genomic architecture, function and evolutionary conservation, suggesting that ZGA follows a conserved molecular logic despite flexible transcriptomic outputs.

## Main text

Across the diversity of animal life, early embryogenesis unfolds through a mixture of deep evolutionary conservation and marked divergence. Animal embryos pass through key developmental milestones, such as fertilization, cleavage, gastrulation and organogenesis. However, these processes often show striking variation across species, reflecting their lineage history, reproductive strategy, and ecology. Another universal yet highly variable developmental milestone is zygotic genome activation (ZGA). Early animal development is initially governed by maternal mRNAs and proteins deposited during oogenesis, until the embryo takes control and begins transcribing its own genome during the maternal-to-zygotic transition (Kojima et al., 2025). ZGA is often considered to start with the transcription of only a few loci, traditionally referred to as the “minor” ZGA wave, which progressively leads to a widespread (“major”) wave of zygotic transcription (Tadros and Lipshitz, 2009). However, recent studies suggest that there is likely a continuum between these two waves (Kojima et al., 2025). Therefore, for simplicity, we hereafter refer to the period of large-scale activation of the zygotic genome as ZGA.

ZGA is essential for subsequent cell fate determination and morphogenesis, and defects in this process lead to developmental arrest (Schulz and Harrison, 2019). Intriguingly, despite its fundamental importance, the timing of ZGA is known to vary extensively across animals. For example, while mice and *Caenorhabditis elegans* initiate large-scale zygotic transcription very early (after one and three cleavage cycles, respectively), other organisms such as frogs, zebrafish and fruit flies do so much later (after approximately 9, 10 and 14 cell cycles, respectively) (Macchietto et al., 2017; Vastenhouw et al., 2019). Moreover, the factors determining these different timings are still unclear, and it is not known to what extent they are shared across animals or specific to particular lineages. Candidate mechanisms include the DNA-to-cytoplasm ratio (related to the nuclear-to-cytoplasmic volume [N/C] ratio), cell cycle lengthening, developmental time, and translation of maternal transcripts, all of which are to some extent interconnected and have been shown to have varying impacts in different model systems (Kojima et al., 2025; Zhou and Heald, 2023).

In addition to the absence of a unified knowledge of the timings and mechanisms behind the ZGA, we lack a systematic understanding of whether some genes are preferentially expressed at ZGA, what roles they play, and how evolutionarily conserved their activation is. A few studies have assessed the conservation of ZGA and maternal genes between closely related species (e.g., mammals (Zhai et al., 2022), flies (Atallah and Lott, 2018) or annelids (Liang et al., 2025)). At larger phylogenetic scales, the earliest transcribed genes at ZGA in mice, zebrafish and fruit flies are, on average, shorter and evolutionary younger, with little overlap between the gene sets activated in these three species (Heyn et al., 2014). Nonetheless, our understanding of ZGA evolution, from its timing to the characteristics of the first zygotic genes expressed, is still greatly limited by the focus on a small set of model organisms, hindering robust evolutionary conclusions.

### Identification of the timing of ZGA across 61 animal species

To obtain a broad phylogenetic view of ZGA and uncover conserved as well as divergent features associated with this process, we assembled a time-course transcriptomic dataset for 61 animal species spanning 13 phyla and more than 700 million years of evolutionary divergence since the common ancestor of ctenophores and mammals (Fig. 1). For most species, this vast dataset consists of eight early embryonic stages, ranging from oocyte/1-cell to post-gastrulation timepoints (Fig. 1), which were generated in-house (23 species, often filling important phylogenetic gaps) or compiled from public sources (38 species). Most samples corresponded to bulk RNA sequencing (RNA-seq), either using protocols for ribosomal RNA depletion (ribo-depletion) or polyA-selection (Table S1). In particular, the majority of the datasets for this study were generated using a single ribo-depletion protocol, following a multi-species pooling strategy to minimize batch effects (see Methods).

**Fig. 1:**
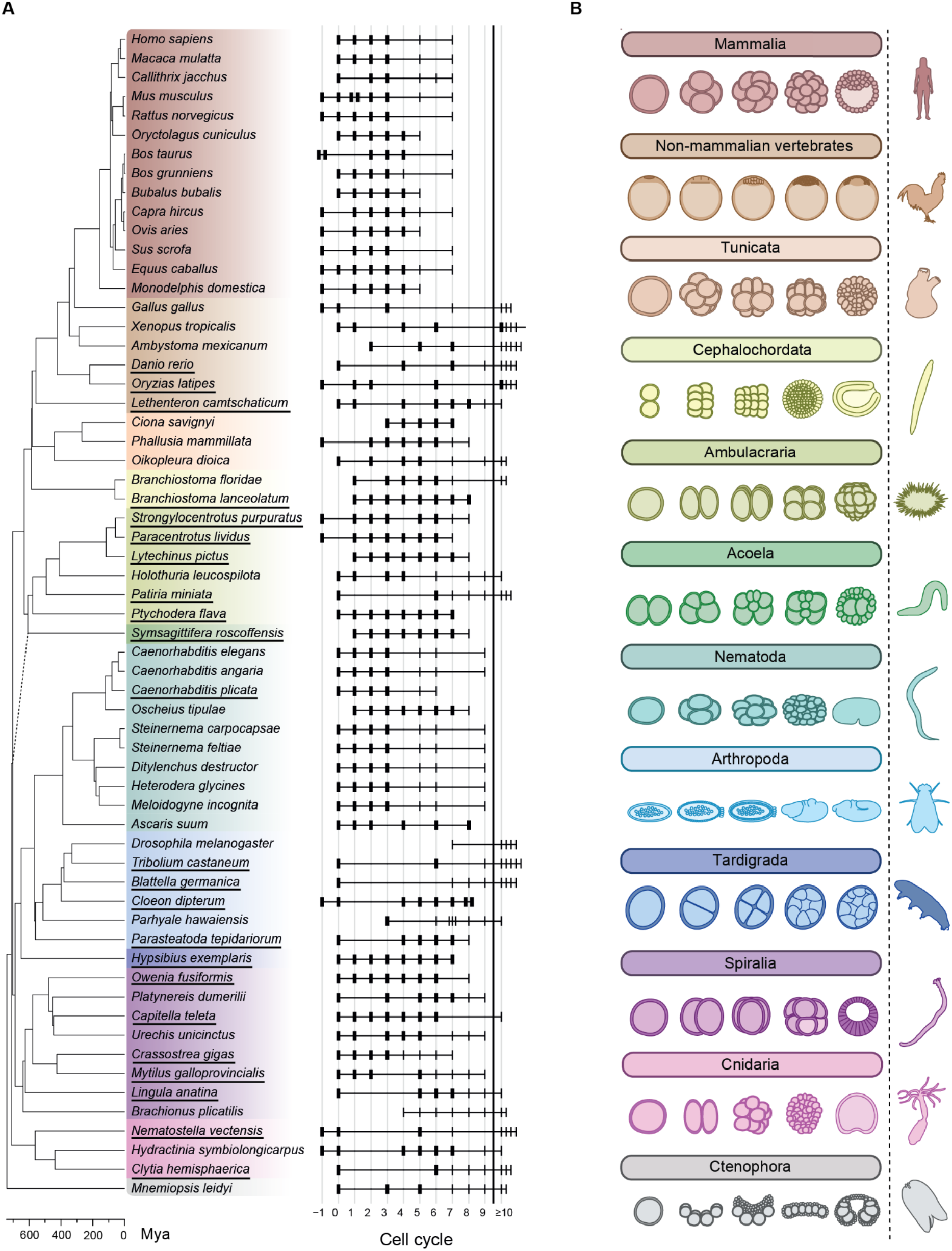
Comparative transcriptomics of early embryogenesis across the animal tree of life. **A**. Phylogenetic framework and dataset overview. For each of the 61 species, early developmental stages are shown on a cell cycle axis; a vertical divider marks the transition to more than 10 mean cell cycles (where absolute cell counts become difficult to estimate, see Methods). A cell cycle value of -1 corresponds to the oocyte, 0 to the 1-cell stage, 1 to the 2-cell stage and so on. Stage markers vary in weight to indicate staging confidence: bold = precise staging; thin = approximate/uncertain (e.g., morula in mammals ≈ 32 cells; some nematode stages reported as 22-44 cells were represented by an average of 32 cells and 5 cycles). Underlined species names denote in-house generated samples, whereas samples for the other species were obtained from public datasets. Note that the non-equivalent taxon groupings are aimed at balancing differences in taxon sampling (especially for mammals and nematodes), as well as reflecting fundamental developmental divergence (see “Phylogenetic framework” in Methods for further details). The dash line indicates the uncertain phylogenetic position of acoels. **B**. Five developmental-stage cartoons for one representative species from each major group (from top to bottom: *Homo sapiens, Gallus gallus, Ciona savignyi, Branchiostoma floridae, Strongylocentrotus purpuratus, Symsagittifera roscoffensis, Caenorhabditis elegans, Drosophila melanogaster, Hypsibius exemplaris, Owenia fusiformis, Nematostella vectensis* and *Mnemiopsis leidyi*). The respective adult organism is shown on the right of the vertical dashed line. Figures are not to scale and are intended only as schematic representations.

Transcriptomic data for all 61 species were processed in a unified manner, mapping the RNA-seq reads against the corresponding genome through STAR, and using standardized genome annotations to extract gene expression levels in transcripts per million (TPM) and intron/exon read counts for each protein-coding gene (see Methods; Table S2, Supplementary Datasets 1,2). First, we reasoned that the earliest sampled stage in each species, often the oocyte or zygote, most closely captures the maternally deposited transcriptome and we therefore defined it as the maternal (MAT) stage. We then employed three complementary metrics to detect the onset of large-scale zygotic transcription in each species (Fig. 2A, Fig. S1A-C and Supplementary Dataset 3): (i) abrupt changes in transcriptome structure captured by principal component analysis (PCA) of the most variable genes, (ii) appearance of strongly upregulated genes that were absent or only weakly expressed at the MAT stage, suggesting activation of zygotic expression, and (iii) increase in the intron/exon read counts ratio, indicative of nascent transcription (Riemondy et al., 2023). These three metrics were scaled within each data type and species so that their maximum value equals 1, and combined to automatically determine the most likely ZGA stage, i.e., the first stage with evidence of substantial new transcription. First, we calibrated this procedure by identifying the most optimal threshold for seven well-characterized model species (Fig. S1D-F; see Methods). Then, we applied this approach to all species in our dataset and performed some minor manual curation (Table S3), generating a comprehensive atlas of the timing of ZGA across animal evolution (Fig. 2B).

**Fig. 2:**
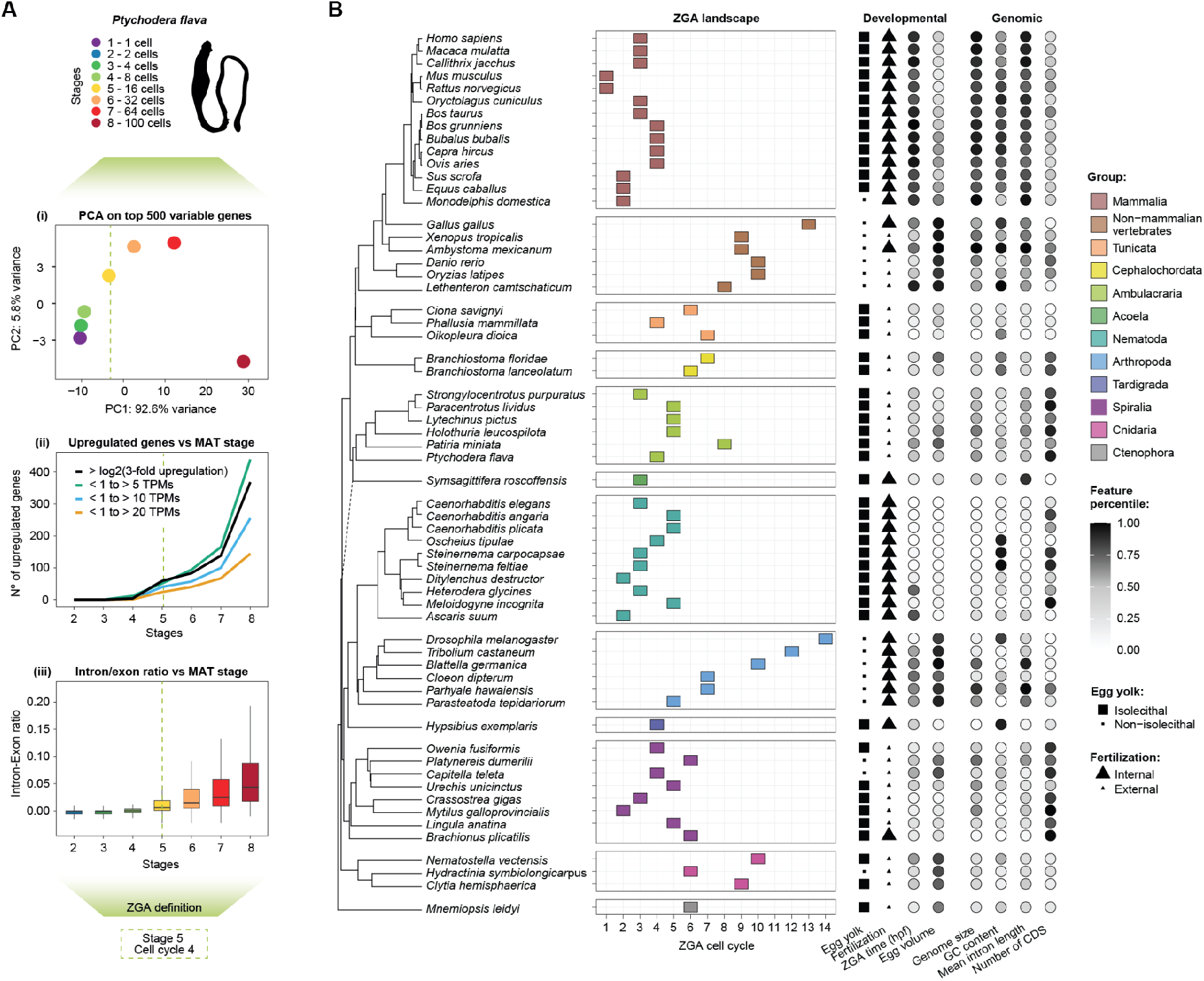
Developmental atlas of ZGA onset across major animal groups. **A**. Overview of the method used to define ZGA onset. A combined metric computed based on three different approaches was employed (example shown for *Ptychodera flava*): (i) a principal component analysis (PCA) of the 500 most variable genes, (ii) expression dynamics of strongly upregulated and newly transcribed genes relative to the first sampled developmental stage (MAT stage), and (iii) increase in the intron/exon read count ratio. **B**. Left: landscape of estimated ZGA onset (measured in cell cycles; x axis) across the animal phylogeny. Right: developmental and genomic features corresponding to each species. hpf: hours post fertilization. The dash line indicates the uncertain phylogenetic position of acoels.

Analysis of the resulting ZGA landscapes revealed considerable variation in the inferred onset of transcription across species, spanning from 1 to 14 cell cycles (Fig. 2B). Some species, including most mammals and nematodes, initiated zygotic transcription within the first two or three cleavages, whereas others, such as many insects and non-mammalian vertebrates, showed much later activation (Fig. 2B). Despite this interspecies variability, ZGA timing tended to be more consistent within clades and exhibited a significant phylogenetic signal (Pagel’s λ = 0.99 and Blomberg’s K = 0.92, *P* ≤ 1e-4; see Methods). However, while ZGA was inferred to occur around cycles 3 to 5 for half of the species (31/61, Fig. S2A), ancestral phylogenetic inference methods could not confidently estimate an ancestral ZGA timing at the root of the phylogeny due to its high variability (estimated at cell cycle 6, with a wide 95% confidence interval = 3.7– 8.3, Fig. S2B).

### The egg DNA-to-cytoplasm ratio is a universal timer of ZGA across animals

Next, we aimed to elucidate what determines this diversity of ZGA onsets. It has been proposed that the timing of ZGA (measured in cell cycles) is inversely associated with the speed of embryonic development (measured in hours) (Kojima et al., 2025). Fast-developing species such as frogs, zebrafish and fruit flies activate their genome after a short amount of time in which multiple quick cell cycles (that omit G-phase) occur. In contrast, slow-developing species such as mammals activate their genome after a longer amount of time in which only a few cell cycles (that include G-phase) take place. These patterns alone would therefore suggest a negative correlation between ZGA cell cycle and developmental timing. However, in our dataset we also observe the opposite trend. Some fast-developing species (e.g., annelids, nematodes, etc.) activate their genome after only a couple of cell cycles (comparable to mammals), while the slow-developing lamprey activates its genome after eight cell cycles and 77 hours of development (Fig. 2B). Consistent with this, we found no significant negative correlation between developmental time at ZGA measured in cell cycles vs. hours (Fig. 3A; adjusted R^2^_OLS_ = 0.017, *P*_PGLS_ = 0.916). In fact, when slow and fast developers were analyzed separately, timing of ZGA in hours and cell cycles showed a mild positive correlation (Fig. 3A; Slow developers: adjusted R^2^_OLS_ = 0.236, *P*_PGLS_ = 0.008; Fast developers: adjusted R^2^_OLS_ = 0.221, *P*_PGLS_ = 0.205).

**Fig. 3:**
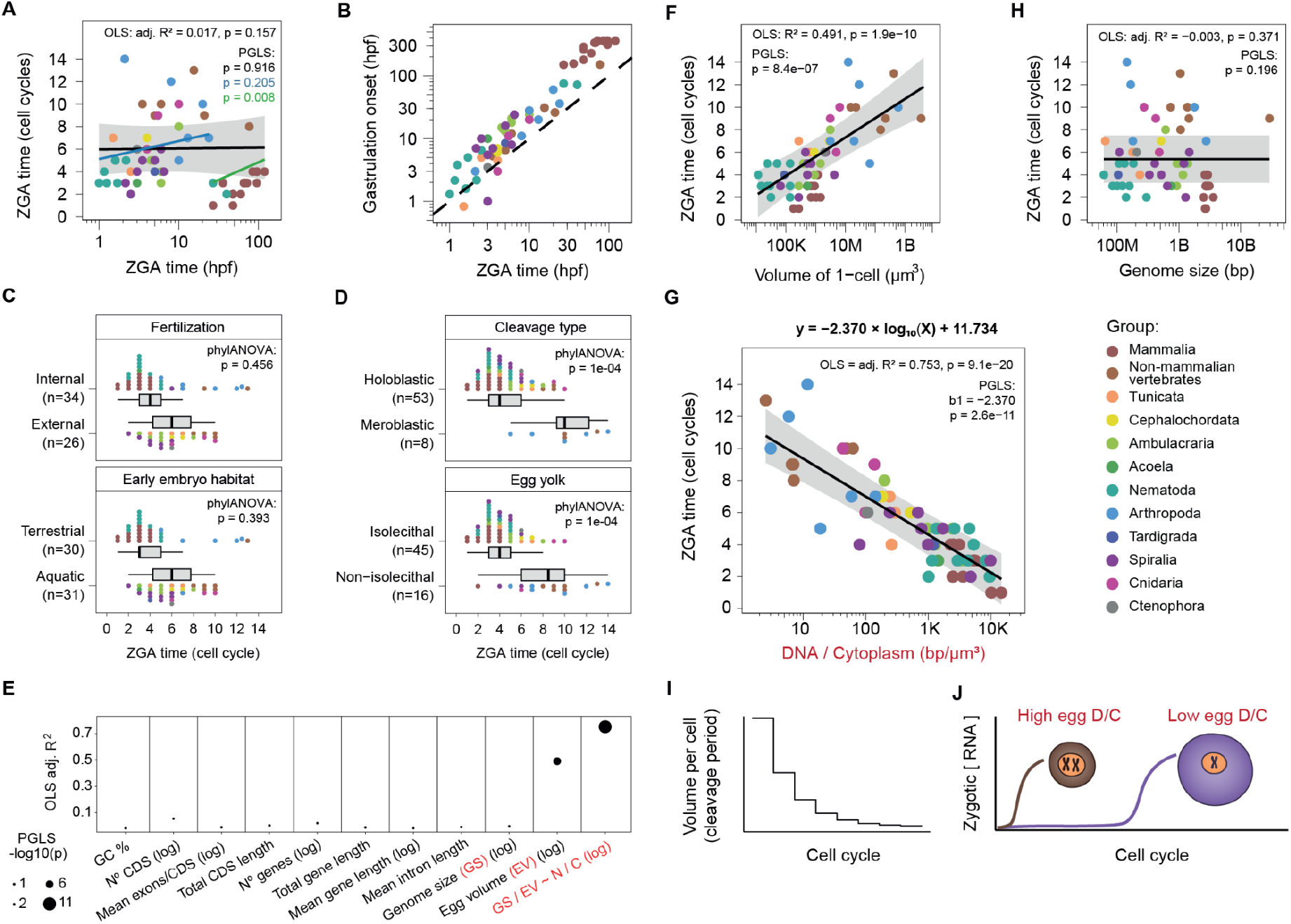
Associations between cell cycle at ZGA onset and multiple biological variables. **A**. Comparison of ZGA onset timing expressed in hours post-fertilization (hpf; x axis) and cell cycle number (y axis). Fitted phylogenetic generalized least squares (PGLS) models are shown for all species combined (black), slow-developing species (green), and fast-developing species (blue). **B**. Comparison of ZGA (x axis) and gastrulation (y axis) onset timings expressed in hpf. **C**. Boxplots representing ZGA onset timings (expressed in cell cycles) across different fertilization strategies (top) and early embryo habitats (bottom). **D**. Boxplots representing ZGA onset timings (expressed in cell cycles) across different cleavage types (top) and egg yolk distribution (bottom). **E**. Predictive power of genomic and developmental features (x axis) for ZGA timing expressed in cell cycles, measured as adjusted R^2^ from ordinary least squares models (R^2^_OLS_) (y axis). For this panel, variables were analyzed either in raw scale or log_10_ scale depending on which gave the higher R^2^_OLS_. Dot size was proportional to -log10(*P*_PGLS_). **F**. Comparison of egg volume in µm^3^ (x axis) and ZGA timing expressed in cell cycles (y axis). **G**. Comparison of egg DNA-to-cytoplasm (D/C) ratio, calculated as genome size (bp) divided by egg volume (µm^3^) (x axis), and ZGA timings expressed in cell cycles (y axis). **H**. Comparison of genome size in bp (x axis) and ZGA timing expressed in cell cycles (y axis). **I**. Schematic of decreasing cell volume per cleavage period. **J**. Model summarizing interspecies differences in ZGA timings as a function of the egg D/C ratio. The circles represent 1-cell embryos from two species. In the brown species, a smaller egg and larger genome produce a higher initial D/C ratio, resulting in earlier ZGA. In the purple species, a larger egg and smaller genome produce a lower initial D/C ratio, resulting in later ZGA. For panels A, F, G and H, black lines indicate PGLS fits estimated with *nlme*. Gray shading shows the confidence interval of the *nlme* PGLS fit. Statistics shown in each panel include OLS R^2^ and the corresponding *P* values for OLS and PGLS. Full statistics are shown in Supplementary Dataset 4. Log-transformed variables refer to log10 transformations.

Another developmental process previously linked with ZGA is the mid-blastula transition (Kojima et al., 2025), characterized by loss of cell cycle asynchrony and changes in cell shape and adhesion that prepare the embryo for the complex morphogenetic processes that will take place during gastrulation. We reasoned that such processes generally require zygotic control and thus must be preceded by ZGA. In line with this, we found that in nearly all species (58/61) ZGA occurs before the onset of gastrulation (Fig. 3B). The exceptions are a small number of fast-developing species (*Oikopleura dioica, Brachionus plicatilis* and *Hydractinia symbiolongicarpus*), in which the earliest detectable signs of gastrulation occur around the same time as ZGA. Overall, these results indicate that gastrulation may impose a developmental constraint on the upper time limit for ZGA, but it is unlikely to directly explain its onset.

In order to identify potentially causal features explaining ZGA onset, we next compiled information on various ecological, developmental and genomic traits for each species (Fig. 2B and Table S4), and assessed their association with ZGA timing measured in cell cycles. Ecological traits such as having internal or external fertilization or an aquatic or terrestrial habitat were not significantly associated with differences in ZGA onset after correcting for phylogenetic structure (Fig. 3C; *P* > 0.05, phylANOVA test). On the other hand, several related developmental features such as the cleavage type (holoblastic vs. meroblastic) and the amount and distribution of the egg’s yolk (isolecithal vs. non-isolecithal) had a significant association with ZGA onset (Fig. 3D; *P* = 1e-04 and *P* = 1e-04, respectively, phylANOVA test). In particular, species with isolecithal eggs (i.e., with little and evenly distributed yolk) and holoblastic cleavage (i.e., early cell divisions that completely divide the egg) exhibited much earlier ZGA compared to species with large amounts of yolk (median ZGA onset: 4 vs. 9-10 cell cycles). We next tested the correlation of a wide range of quantitative traits with ZGA onset (Fig. 3E). Most of these traits were not significantly correlated (Fig. 3E, Fig. S3A,B and Supplementary Dataset 4), with the exception of egg volume, which showed a robust positive correlation with ZGA onset (Fig. 3F, adjusted R^2^_OLS_ = 0.491, *P*_PGLS_ = 8.4e-7). Since egg size primarily reflects cytoplasmic volume, this observation suggested that the amount of cytoplasm relative to nuclear content may be a key determinant of ZGA onset (Kojima et al., 2025). This is consistent with the long-standing model in which the N/C ratio regulates the onset of zygotic transcription. We therefore tested whether a composite trait capturing both the egg’s DNA content and cytoplasmic volume, serving as a proxy for its N/C ratio, could better explain the observed variation in ZGA timing. As predicted, the logarithm of the ratio between genome size and egg volume showed a striking negative correlation with ZGA timing (Fig. 3G, adjusted R^2^_OLS_ = 0.753, *P*_PGLS_ = 2.6e-11). This correlation was stronger than egg volume alone, even despite the fact that genome size and ZGA onset were not significantly correlated (Fig. 3H, adjusted R^2^_OLS_ = -0.003, *P*_PGLS_ = 0.196). Importantly, the correlation between egg DNA-to-cytoplasm ratio and ZGA onset remained robust and highly significant when only specific subsets of species were used, including those sequenced through ribo-depletion or polyA-selection (Fig. S4A,B), those generated for this study or compiled from public sources (Fig. S4C,D), or for slow and fast developers separately (Fig. S4E,F). Altogether, these results strongly suggest that the egg DNA-to-cytoplasm ratio, and its progressive increase with each cleavage cycle, act as a universal timer of ZGA across animals (Fig. 3I,J). This also allowed us to develop a simple predictor of the ZGA time (in cell cycles) for any species of interest, provided its genome size and egg volume are known: ZGA time (cell cycle) = -2.370 * (log_10_(GS/EV)) + 11.734, where GS corresponds to genome size (in bp) and EV to egg volume (in um^3^).

### Genes transcribed at ZGA share distinctive molecular and functional features

We next investigated the characteristics of the genes expressed at ZGA. Given the large phylogenetic variability of ZGA timing, number of protein-coding genes, and type and coverage of RNA-seq data, we implemented a simple score (ZGA_score_) to systematically compare zygotic genes across animals. This score ranks all genes within each species according to the extent of their activation at ZGA, comparing each gene’s expression at ZGA and the subsequent stage to its expression at the MAT and pre-ZGA stages (see Methods, Supplementary Dataset 5). For each species, the top 500 genes by ZGA_score_ were designated as the set of “ZGA genes”. For comparison, we also defined a set of maternally deposited genes in each species as those with high expression (TPM ≥ 50) at the MAT stage (“MAT genes”). As expected, ZGA genes showed a sharp rise in expression at the ZGA stage across species, while MAT genes exhibited a mild decline (Fig. 4A and Fig. S5A,B). We then set out to characterize the ZGA and MAT gene sets from a molecular and functional perspective, to determine whether each set displays conserved features across animals.

**Fig. 4:**
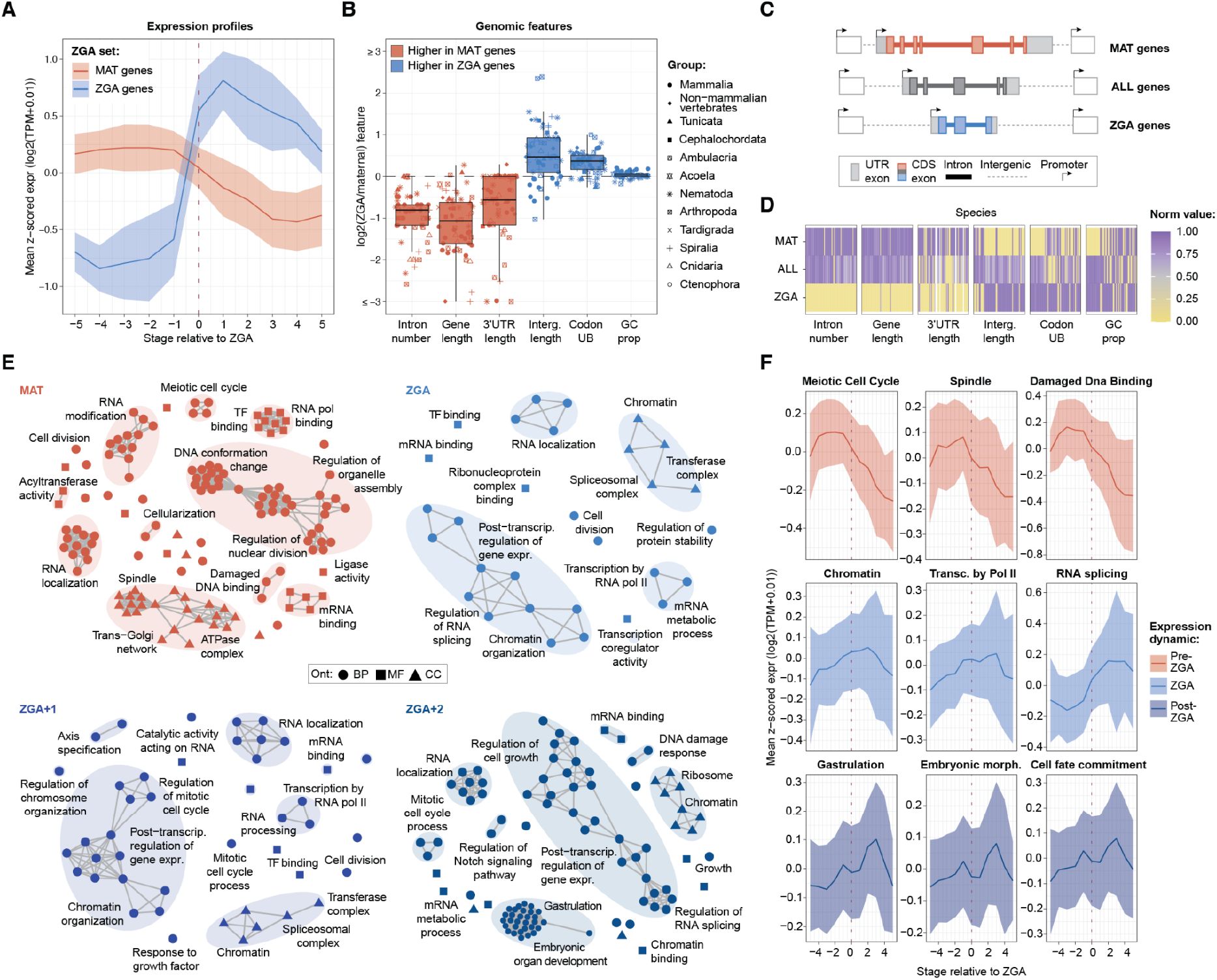
Comparison of molecular features between ZGA and MAT genes. **A**. Summarized expression across developmental stages for ZGA (blue) and MAT (red) gene sets across all species. Expression values represented as log_2_(TPM + 0.01) were z-scored across stages per gene and species, averaged within gene sets per species, and then summarized across species (see Methods). Thick lines show the across-species mean and shaded regions indicate ±1 standard deviation. Stages were aligned relative to the ZGA stage (0). **B**. Boxplots showing within-species comparison of genomic features (x axis) between MAT and ZGA genes represented as log_2_ fold change (y axis). Dot shapes represent phylogenetic groups, while boxplot colors indicate whether a given feature is overall higher for MAT genes (red) or ZGA genes (blue). **C**. Schematic representation of genomic features for MAT, ZGA and all genes. **D**. Heatmap showing within-species comparison of genomic features (x axis) between MAT genes, ZGA genes, and all genes with expression ≥ 1 TPM in at least one stage. Feature values within each species were normalized by the maximum observed value and the lowest and highest values set to 0 and 1, respectively. See Fig. S5C for the full version. **E**. Networks of significant GO categories from GSEA run on percentiles calculated on decreasing (i) expression at MAT stages, (ii) ZGA scores, (iii) ZGA_+1_ scores and (iv) ZGA_+2_ scores. The networks were built through *Revigo v1*.*8*.*2* (Supek et al., 2011) requiring output lists of medium (0.7; ZGA, ZGA+1) or small (0.5, MAT, ZGA+2) sizes, using *SimRel* as similarity measures and removing obsolete terms. Nodes represent GO categories, with shapes indicating the ontology and edges connecting terms with similarity ≥ 0.2. Clusters are highlighted by shaded ellipses and labelled by at least one representative term. A full list of significant GSEA results and relevant statistics is available in Table S6 and depicted in Fig. S6. **F**. Summarized expression across developmental stages for representative significant categories collectively peaking before ZGA (red), at ZGA (light blue) and after ZGA (dark blue). The thick line represents the across-species mean, and the shaded region indicates ±1 standard deviation. Expression values were transformed as described for panel A. Abbreviations: Codon UB: codon usage bias, GC prop: GC proportion, Ont: ontology, BP: biological process, MF: molecular function, CC: cellular component, Transc. by pol II: transcription by RNA pol II, RNA splicing: regulation of RNA splicing, Embryonic morph: embryonic morphogenesis.

Comparative analyses revealed highly consistent genomic differences between MAT and ZGA genes for various tested features (see Methods and Table S5). First, ZGA genes were shorter and contained fewer introns (Fig. 4B), architectural features that enable rapid transcription during short cleavage cycles (Kojima et al., 2025). Second, they tended to have shorter 3′ UTRs and higher codon usage bias (Fig. 4B), suggesting simpler and less efficient regulation of their translation and mRNA stability as opposed to MAT genes (Bazzini et al., 2016; Mishima and Tomari, 2016). Third, with the exception of mammals, they displayed larger intergenic regions (Fig. 4B), supporting more complex transcriptional regulation (Marlétaz et al., 2018; Nelson et al., 2004). Finally, ZGA genes generally showed higher GC content than MAT genes (48/61 species, 79%), even if the magnitude of these differences was smaller compared to the other analyzed features. Comparisons with the genomic median (see Methods) allowed for the quantification of the deviation of each feature for ZGA and MAT genes and hinted at their archetypal profiles across animals (Fig. 4C,D and Fig. S5C).

We then searched for functions shared by genes activated during ZGA. To perform an unbiased comparison across species and developmental stages, we first generated two additional scores for each gene and species, which are equivalent to the ZGA_score_ but measure the extent of activation either one or two stages after ZGA in our time courses (ZGA+1_score_ and ZGA+2_score_, respectively) (Supplementary Dataset 5; see Methods). For the MAT stage, we simply ranked genes according to their expression level within each species, which reflects their level of maternal deposition. Next, we compiled standardized Gene Ontology (GO) annotations of all protein-coding genes in each species (see Methods), and ran Gene Set Enrichment Analyses (GSEA) for each species and focus stage (MAT, ZGA, ZGA+1 and ZGA+2), using each ranked gene list as input (Supplementary Dataset 6). This analysis revealed various gene functions that are consistently enriched across the phylogeny for each stage (Fig. 4E, Fig. S6 and Table S6). For the MAT stages, commonly enriched GO terms included DNA damage response, microtubule-based processes, mitotic cell cycle, RNA localization, and post-transcriptional regulation of gene expression. For ZGA and ZGA+1 stages, enriched GO terms were mainly related with RNA processing and gene expression, including chromatin, transcription by RNA polymerase II, mRNA binding, and splicing, as well as a few terms associated with early embryogenesis. In contrast, the ZGA+2 stages showed enrichments that predominantly reflect post-cleavage developmental programs. Such terms could be grouped into some main categories: patterning and axis formation, morphogenesis and gastrulation, organogenesis across diverse systems, and structural and signaling processes. These common functional developmental trajectories across species could be further observed by aggregating across all species relative gene expression dynamics over developmental time within selected GO categories (Fig. 4F). Altogether, these enrichments reflect a conserved functional switch from maternal control of cell division to first zygotic reprogramming of transcriptional and chromatin landscapes, and ultimately to embryogenesis and organogenesis, consistent with the temporal needs of early developmental progression.

### Zygotic genes show low genomic and regulatory conservation across animals

Although our results revealed that similar gene functions are activated at the ZGA in many species, previous studies suggested little conservation at the gene level among three model organisms (Heyn et al., 2014). Therefore, we assessed the evolutionary conservation of ZGA genes, both in terms of ortholog presence/absence (genomic) and the activation of these orthologs at ZGA (regulatory) in other species. For this purpose, we first inferred orthology relationships among all protein-coding genes from the 61 species in the form of gene orthogroups (Supplementary Dataset 7). For each species, we measured the genomic conservation of the ZGA and MAT gene sets in all the other species, relative to a background comprising all protein-coding genes minus these two sets (see scheme in Fig. S7A). We considered a query gene to be genomically conserved in a target species when at least one of its orthologs was present in that species. For all species, MAT genes exhibited higher genomic conservation than the background (Fig. 5A and Fig. S7B). In stark contrast, ZGA genes showed globally lower conservation than the background and always lower conservation than MAT genes (Fig. 5A and Fig. S7B,C). Similar results, albeit with noisier patterns, were obtained using a phylostratigraphic approach to define old and new genes within each species (see Methods), which revealed that ZGA genes were overall newer than MAT genes (Fig. S7D). Consistent with these findings, the fraction of ZGA genes present in any orthogroup was much lower than that of MAT genes across all species (Fig. S8A).

**Fig. 5:**
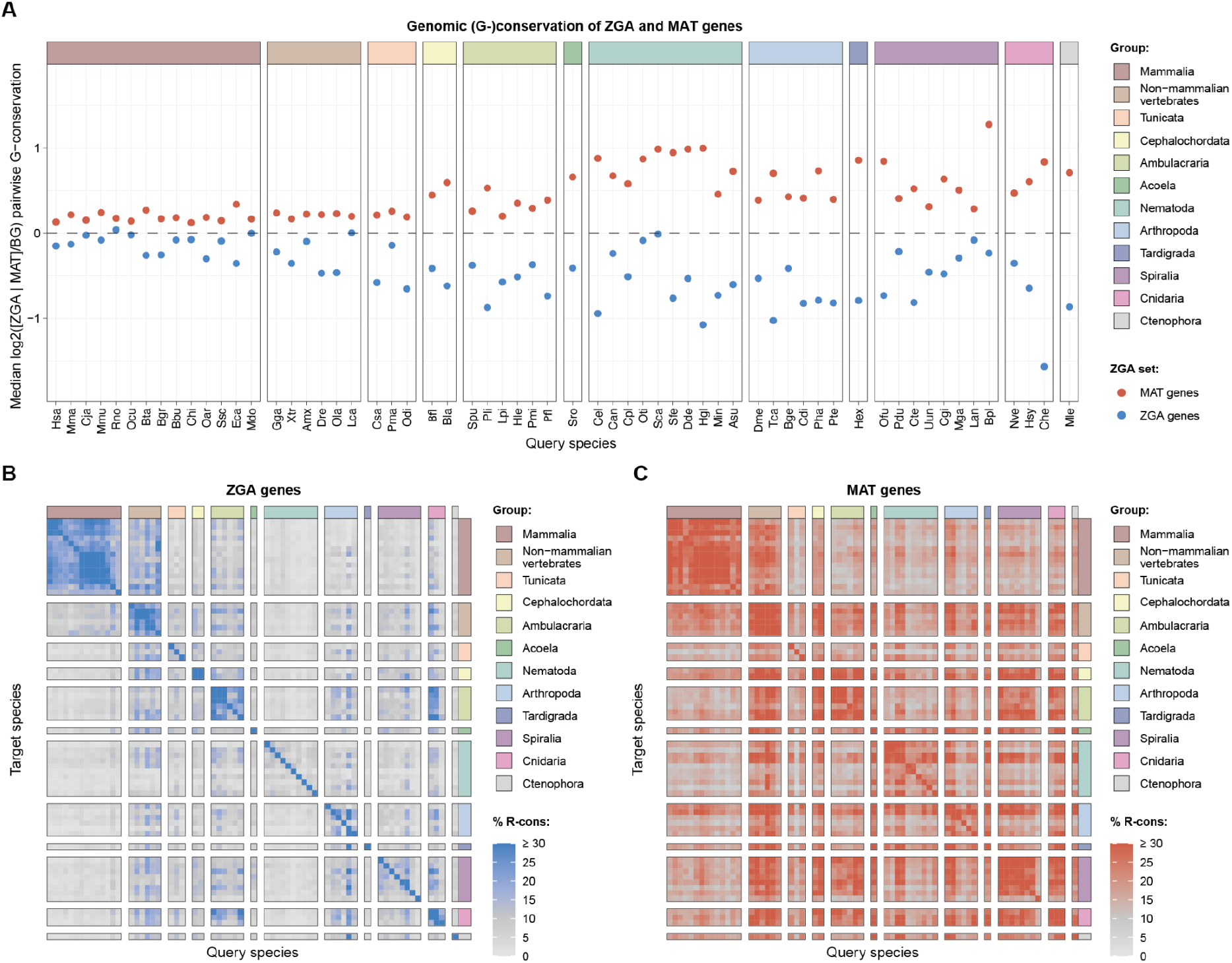
Evolutionary conservation of ZGA and MAT genes. **A**. Genomic (G-)conservation of the ZGA and MAT gene sets in each species relative to its genomic background. A query gene is considered G-conserved in a target species if that species contains at least one gene in the same orthogroup. The plotted value (y axis) for each query species (x axis) represents the median log_2_ G-conservation ratio (ZGA or MAT vs the relative background) in all target species (full distributions in Fig. S7B,C). Positive / negative values on the y axis indicate more / less G-conservation than the respective backgrounds. For each query species, the background set is composed of its protein-coding genes depleted of the tested sets. **B**,**C**. Percentage of ZGA (B) or MAT_top_ (C) genes from each query species (x axis) with regulatory conservation (R-conservation) into each target species (y axis). A query ZGA/MAT_top_ gene is considered R-conserved in a target species if that species contains at least one ZGA/MAT_top_ gene in the same gene orthogroup. Note that each ZGA and MAT_top_ set consists of 500 genes.

Next, we assessed whether the orthologs of ZGA genes in a given species tended to also be strongly activated at ZGA in the other species (regulatory conservation). For this purpose, we performed pairwise comparisons among all species by computing the fraction of ZGA genes in the query species that had at least one ZGA ortholog in the target species (see Methods). This revealed that ZGA genes mainly exhibit regulatory conservation within certain groups (e.g., mammals, ambulacraria or cnidarians), with minimal conservation across them (Fig. 5B). In contrast, a comparable analysis for the top maternally deposited genes in each species revealed higher fractions of conserved expression profiles across the phylogeny (Fig. 5C and Fig. S8B; see Methods).

These observations prompted us to define both group-specific and universally-shared ZGA and MAT gene orthogroups (see Methods). On one hand, consistent with the previous analysis, ZGA genes were associated with more group-specific orthogroups than MAT genes (Fig. S8C-F). Notably, these group-specific ZGA orthogroups generally had no detected homologs in any of the other animal groups (Fig. S8E), suggesting a strong association between gene emergence and the acquisition of ZGA regulation. For instance, mammalian-specific ZGA orthogroups were enriched for GO categories related to response to stimuli, transcriptional activation and RNA biosynthesis (Fig. S9). One such orthogroup included the human *TRIM43* and its homologs, a family of ubiquitin-ligating proteins that emerged in the mammalian ancestor and representing well-known markers of mammalian ZGA (Stanghellini et al., 2009; Taubenschmid-Stowers et al., 2022). On the other hand, ZGA genes were associated with far fewer universally shared orthogroups compared to MAT genes (Fig. S8C-F). In fact, only one orthogroup was consistently activated at ZGA across all groups (Fig. S8D,E), corresponding to the human FET genes (FUS, EWSR1 and TAF15) and their homologs across animals. FET genes are RNA binding proteins that mediate co-transcriptional RNA processing, and their perturbation has been linked to splicing defects affecting gastrulation (*Fus* in the clawed frog (Dichmann and Harland, 2012)), mitotic defects (*ewsr1a* and *ewsr1b* in zebrafish (Azuma et al., 2007)) and contributes to differential post-ZGA RNA binding (*caz* in fruit fly (Sysoev et al., 2016)). In contrast, we detected 522 broadly shared MAT gene orthogroups (Fig. S8D,Ff), with functional enrichments reflecting those revealed by GSEA at the MAT stages (figs. S6 and S9). Examples of such genes include the membrane-associated cytoskeletal proteins Ezrin and Radixin, whose combined knockdown in mouse significantly decreases fertilization success (Cohen et al., 2022). In summary, these results show that ZGA genes tend to be group- or even species-specific, whereas a common set of MAT genes mainly involved in regulation of cell cycle and RNA processing are universally deposited as mRNAs in eggs during oogenesis.

## Discussion

In this study, we took a comparative approach to systematically investigate ZGA across animals. By compiling a vast transcriptomic dataset of early developmental stages for 61 species, we generated a comprehensive atlas of the timing of ZGA. We found that large-scale activation of the zygotic genomes can occur at a wide range of cell cycles after fertilization (from 1 to 14), and it is highly variable across animal groups. Despite this large evolutionary divergence, we found the ratio between the egg’s DNA and cytoplasmic volume to likely act as a universal timer of ZGA, implying that the zygotic gene transcription starts once a certain cellular DNA-to-cytoplasm ratio is reached. Moreover, we also revealed multiple genomic, functional and evolutionary features broadly associated with genes activated at ZGA. This points to a common molecular logic for this essential developmental process across animals, and opens the possibility that similar rules operate across species despite the overall lack of overlap between individual zygotic genes.

We uncovered a negative correlation between the number of cell cycles at which the ZGA is detected in a given species and a proxy for the egg N/C ratio based on genome size and egg volume. Importantly, this correlation was strong and robust despite some potential caveats of our dataset, such as the wide variation in actual times for equivalent stages (e.g., from < 20 minutes for the first cleavage in the tunicate *O. dioica* (Wang et al., 2025) to 20-24 hours in mammals), the diversity of RNA-seq data types (e.g., ribo-depletion vs. polyA-selected), the lack of nascent RNA-seq, and the bulk nature of the transcriptomes (specially problematic for species with late activation and with highly varying blastomere sizes). Another consideration is related to the difficulty of estimating the fraction of “active” cytoplasmic volume (i.e., containing the maternal mRNAs and proteins) relative to that of “inactive” yolk. Thus, it is plausible that the association between egg DNA-to-cytoplasm ratio and ZGA timing is even stronger than reported here. The N/C ratio has long been proposed as a key determinant of ZGA onset, with a negative association reported in a few model animal species (Kojima et al., 2025), but also a surprisingly contrasting pattern emerging from a recent study in brown algae (Bogaert et al., 2026). Our findings not only uncover a widespread negative association between DNA-to-cytoplasm and ZGA onset across dozens of animal species, but also strongly support its role as a more universal timer. In fact, incorporating the data available for two plant species (Arabidopsis (Kao and Nodine, 2019) and maize (Chen et al., 2017)) reveals a pattern highly consistent with that observed in animals and further strengthens the overall negative correlation observed here (Fig. S10, adjusted R^2^_OLS_ = 0.772, *P*_PGLS_ = 2.1e-18), suggesting that this relationship may extend across other multicellular eukaryotes.

How can the egg DNA-to-cytoplasm ratio act as a ZGA timer? Since during early animal cleavages the DNA amount doubles but the overall embryo volume remains essentially constant, every cell cycle exponentially increases the proportion between the amount of genomic DNA and the cytoplasmic material (Fig. 3I), in line with the negative linear relationship we observed between the logarithm of this ratio in the egg and the cycle of ZGA. This suggests that ZGA may occur when a maternally deposited factor acting on the DNA is titrated enough after various rounds of genomic replication (Kojima et al., 2025). Consistently, triploid and haploid cybrid frog embryos respectively displayed earlier and later ZGA compared to diploid embryos (Jukam et al., 2021), and haploid honeybee males activate their genome later than diploid females (Wang et al., 2021). Histones are long-standing candidates as such factors whose titration or depletion leads to ZGA onset. In frog, titration of histones was shown to help set ZGA timings (Amodeo et al., 2015). In zebrafish, ZGA timing depends on the balance between nucleosome-forming histones and transcription factors, with declining availability of non-DNA-bound histones allowing transcription factors to compete more effectively for genomic binding (Joseph et al., 2017). However, the DNA-to-cytoplasm ratio is also tightly connected with cycle length, which has also been shown to directly influence ZGA timing (Zhou and Heald, 2023). This effect can be mediated by inhibition of cell cycle regulators by histones, as seen in fruitfly (Shindo and Amodeo, 2021), or directly through the dilution of replication factors, which also act on the DNA and lead to slower cell cycles, as reported in frogs (Collart et al., 2013). Therefore, the DNA-to-cytoplasm ratio could cause the lengthening of the cell cycle, which in turns triggers ZGA. Moreover, it is possible, or even likely, that a combination of both direct and indirect determinants operates to different extents in each species. Whatever the underlying mechanism(s), however, our comparative atlas of ZGA across animals provides strong support for the idea that the egg DNA-to-cytoplasm ratio acts as a universal timer of genome activation across animals, and reveals that this fundamental developmental transition follows a conserved functional logic despite extensive evolutionary turnover of the underlying genes.

## Materials and Methods

### Phylogenetic framework

The species included in our dataset (and the respective scientific acronyms) are the following: *Homo sapiens (Hsa), Macaca mulatta (Mmu), Callithrix jacchus (Cja), Mus musculus (Mmu), Rattus norvegicus (Rno), Oryctolagus cuniculus (Ocu), Bos taurus (Bta), Bos grunniens (Bgr), Bubalus bubalis (Bbu), Capra hircus (Chi), Ovis aries (Oar), Sus scrofa (Ssc), Equus caballus (Eca), Monodelphis domestica (Mdo), Gallus gallus (Gga), Xenopus tropicalis (Xtr), Ambystoma mexicanum (Ame), Danio rerio (Dre), Oryzias latipes (Ola), Lethenteron camtschaticum (Lca), Ciona savignyi (Csa), Phallusia mammillata (Pma), Oikopleura dioica (Odi), Branchiostoma floridae (Bfl), Branchiostoma lanceolatum (Bla), Strongylocentrotus purpuratus (Spu), Paracentrotus lividus (Pli), Lytechinus pictus (Lpi), Holothuria leucospilota (Hle), Patiria miniata (Pmi), Ptychodera flava (Pfl), Symsagittifera roscoffensis (Sro), Caenorhabditis elegans (Cel), Caenorhabditis angaria (Can), Caenorhabditis plicata (Cpl), Oscheius tipulae (Oti), Steinernema carpocapsae (Sca), Steinernema feltiae (Sfe), Ditylenchus destructor (Dde), Heterodera glycines (Hgl), Meloidogyne incognita (Min), Ascaris suum (Asu), Drosophila melanogaster (Dme), Tribolium castaneum (Tca), Blattella germanica (Bge), Cloeon dipterum (Cdi), Parhyale hawaiensis (Pha), Parasteatoda tepidariorum (Pte), Hypsibius exemplaris (Hex), Owenia fusiformis (Ofu), Platynereis dumerilii (Pdu), Capitella teleta (Cte), Urechis unicinctus (Uun), Crassostrea gigas (Cgi), Mytilus galloprovincialis (Mga), Lingula anatina (Lan), Brachionus plicatilis (Bpl), Nematostella vectensis (Nve), Hydractinia symbiolongicarpus (Hsy), Clytia hemisphaerica (Che), Mnemiopsis leidyi (Mle)*. We assembled them into a phylogenetically calibrated tree (introduced in Fig. 1A) primarily through the *TimeTree* database (Kumar et al., 2017). When one of our species was not available in *TimeTree*, we replaced it with the closest available congener or sister lineage, while, for taxa completely missing from the database, we reviewed the primary literature to correctly place them and assign calibration dates.

Note that, throughout the manuscript, we have used non-equivalent taxon groupings at the rank level (see Fig. 1A), and included the paraphyletic group of “non-mammalian vertebrates”, with the goal of balancing differences in taxon sampling (especially for mammals and nematodes), as well as reflecting fundamental developmental divergence. For instance, members of Chordata (such as humans and fish) exhibit profound disparities in early ontogeny, including yolk volume, cleavage symmetry, and blastula architecture, whereas the Spiralian phyla (such as Mollusca and Annelida) maintain a highly conserved and homologous spiral cleavage program despite their disparate adult morphologies.

### Sample collection and RNA-seq data generation

Public samples were obtained from the following National Center for Biotechnology Information (NCBI) BioProjects: PRJNA1031543, PRJNA1071214, PRJNA1112135, PRJNA1126096, PRJNA1190105, PRJNA198514, PRJNA228235, PRJNA275011, PRJNA312389, PRJNA341580, PRJNA342320, PRJNA343030, PRJNA344880, PRJNA384004, PRJNA394029, PRJNA453988, PRJNA543590, PRJNA579328, PRJNA646061, PRJNA686960, PRJNA688000, PRJNA783716, PRJNA787530, PRJNA821380, PRJNA880794, PRJNA882848, PRJNA933618, PRJNA970726 and PRJNA990458, as well as from the China National GeneBank DataBase (CNGBdb) project CNP0000891

In-house sampling of early developmental stages was performed for 23 animal species, following the protocols described in detail in the Supplementary Methods. To prepare the libraries for Illumina sequencing, given the large phylogenetic divergence of most species, we generated pools with several species together to increase final RNA quantity and reduce batch effects, as described in Table S1. Pools of RNA or RNA from individual species were then processed using ribo-depletion with the TruSeq Stranded Total RNA Kit (Illumina) following the manufacturer’s protocol. For *Caenorhabditis plicata* and *Lingula anatina*, we used poly(A) selection library construction, following the manufacturer’s protocol. Libraries were sequenced on an Illumina NovaSeq 6000 to generate 150 bp paired-end reads. All in-house generated samples were registered under the NCBI BioProject PRJNA1402383. Detailed information, including SRA IDs, for all in-house and public samples can be found in Table S1.

### Bulk RNA-seq mapping and gene expression quantification

We aligned RNA-seq reads to the appropriate reference genomes using STAR *v2*.*7*.*11a* (Dobin et al., 2013), and we built genome indices from the species genome FASTA and gene annotation (GTF) files. We ran alignments in two-pass mode with permissive multimapping to retain repetitive signal using *--winAnchorMultimapNmax 200*, and we sorted the output BAMs by coordinate. We then selected only protein-coding genes (i.e., those with CDS entries) and quantified paired-end RNA-seq fragments at the gene level using *featureCounts v2*.*0*.*2* (Liao et al., 2014). We computed the coding exonic signal by counting fragments over CDS features with fragment-level counting (*-p* option) and a read mapping quality filter of *MAPQ* ≥ *30*. In parallel, we obtained the intronic signal by counting fragments over intron features with the same *MAPQ* ≥ *30* cutoff and requiring ≥ 30 bp overlap into introns. For each developmental stage, we then aggregated counts across biological replicates, when present, to produce a single stage-level quantification per gene, with exon and intron counts computed in parallel. Finally, we converted counts to length-scaled TPMs, removed genes with extreme artifacts (i.e., TPM > 100,000), and renormalized each sample to 10^6^ TPMs.

### Definition of nomenclature for key developmental stages

For each species, the first sampled stage (which generally corresponds to the oocyte or zygote stage) was designated as the maternal (MAT) stage under the assumption that it most closely represents the maternally deposited transcriptomic landscape. The stage corresponding to the onset of zygotic genome activation was designated ZGA, the immediately preceding stage ZGA_pre_, and the two subsequent stages, when available, ZGA_+1_ and ZGA_+2_. This nomenclature was used consistently throughout the manuscript.

### Detection of ZGA timing from bulk RNA-seq data

To consistently detect ZGA onset separately in each species, we used three complementary metrics describing transcriptomic differences between developmental stages that were later integrated into a single composite measurement: a principal component analysis (PCA), comparison of newly transcribed and upregulated genes, and comparison of intron/exon (I/E) read count ratios.

#### Principal component analysis (PCA)

First, we performed a PCA on the stage-level expression matrix derived from the gene TPM values after variance stabilization with the *rlog* transformation in *DESeq2* (design ∼1) (Love et al., 2014), restricting the analysis to the 500 most variable genes across the time course. In most species, stage projections onto the PC1-PC2 plane followed a horseshoe-shaped trajectory that recapitulated developmental progression. Because ZGA is expected to correspond to the first large transcriptomic remodeling across early stages, we reasoned that the first two principal components should separate pre-ZGA from post-ZGA samples, with this key transition marked by a first large distance between consecutive stages. To quantify this transition, we calculated the normalized variance-weighted Euclidean distance in the PC1-PC2 plane between each stage and the immediately preceding one, with PC1 and PC2 weighted by their respective proportions of explained variance.

#### Newly transcribed and upregulated genes

Second, we quantified global transcriptional activation by identifying newly transcribed and strongly upregulated genes at each developmental stage relative to the MAT stage. Genes were classified as newly transcribed at stage *i* when their expression at MAT was < 1 TPM and their expression at stage *i* exceeded 5 TPMs. To capture broader activation beyond strict *de novo* transcription, we also included genes showing a log2 fold change > 3 relative to the MAT stage, after excluding genes with MAT stage expression lower than 10 TPMs. For each stage, we then calculated the union of newly transcribed genes and genes strongly upregulated relative to the MAT stage, providing a broader readout of ZGA. The ZGA stage is expected to correspond to the first sharp increase in transcriptional activation relative to earlier stages.

#### Intron/exon (I/E) read count ratios

Third, we analysed intron-to-exon (I/E) read count ratios across developmental stages, reasoning that this metric should increase at ZGA onset as newly synthesized pre-mRNAs contribute intronic signal before splicing is completed, while maternal transcripts are deposited as fully processed. We restricted this analysis to intron-containing genes with exonic counts ≥ 50 at every stage. For each gene and stage, we calculated the I/E ratio as the number of intronic reads divided by the number of exonic reads. To enrich the expected I/E peak at ZGA, we applied maternal masking filters. This was particularly important because (i) many of the analysed species are non-model organisms with potentially suboptimal genome annotations, (ii) abundant maternally deposited transcripts can generate residual intronic signal at the MAT stage, and (iii) low sequencing depth in some species yields very few reads mapping to exons and introns. Specifically, we retained only genes with a MAT-stage I/E ratio ≤ 0.05 and in any case no greater than 1.5 times the ratio at the last stage. We then quantified stage-specific changes relative to MAT as I/E Δ = ratio*i* − ratio_MAT_ . To reduce the influence of extreme values, we removed the upper and lower 1% of Δ values. Stage-wise I/E Δ distributions were summarized using boxplots. Under this framework, the ZGA stage is expected to correspond to the first sharp increase in I/E read count ratio relative to preceding stages. Plots and summary tables for each species for these three metrics are provided in Supplementary Dataset 3.

#### Integration of the three metrics and inference of ZGA onset

To automate ZGA onset calling across species, we integrated the three metrics above into a composite score. For each species, values for each metric were scaled to the species-specific maximum, yielding normalized values between 0 and 1. We then calculated, for each developmental stage, the mean of the three normalized metrics, and derived the average normalized delta (ZGA Δ) as the change in this composite score between consecutive stages. The predicted ZGA stage was defined as the first stage at which ZGA Δ exceeded a fixed threshold. To select this threshold, we scanned values from 0.001 to 1 in steps of 0.001 using seven model species in which the major wave of ZGA had been well established and extensively characterized (*Homo sapiens, Mus musculus, Danio rerio, Oryzias latipes, Xenopus tropicalis, Caenorhabditis elegans* and *Drosophila melanogaster*). Based on this analysis, we selected 0.1 as the value showing maximal agreement with their previously reported ZGA stages (Fig. S1F). We then applied the same approach with the threshold of 0.1 to detect the ZGA stage in all other species in our dataset. All automatic predictions were subsequently reviewed by visual inspection of the underlying metric profiles, raw metric trajectories and stage-to-stage deltas (Supplementary Dataset 3). This led to targeted manual curation of the detected ZGA stage in seven species showing more complex dynamics, typically owing to noise in one of the three metrics or to weaker secondary ZGA waves (see Table S3 for details).

### Estimation of ZGA timing in cell cycle and absolute time units

To enable cross-species comparison of ZGA, we recorded ZGA onset in both cell cycle units and absolute developmental time (hours). In species with an early ZGA onset and synchronous early cleavages, the cell cycle at ZGA could be inferred readily by cell counting (Fig. 1A). However, in species in which the ZGA occurred after asynchronous cleavages (e.g., nematodes) or at later stages (which are normally asynchronous for all species), we determined the ZGA onset in cycles by inferring the mean cell cycle from the total number of cells in the embryo. For example, a stage with approximately 300 cells was assigned a mean cell cycle of 8 (2^8 = 256). For simplicity, the mean cell cycle is referred to throughout the text as “cell cycle.”

We also used ZGA timing in absolute developmental time in hours, which is simply recorded upon sample collection. Although this was reported for most species as hours post fertilization (hpf), some of the public sources used alternative reference points, including hours post copula, hours after ootheca formation, or hours after egg extrusion. For simplicity, we used “hpf” in the main text as a comparative label for absolute time in hours, while the original timing reference for each species is provided in Table S4.

### Classification of fast and slow developers

Species were assigned to fast- or slow-developing categories using ZGA timing in hpf as a proxy for early developmental rate. This classification was based on the distribution shown in Fig. 3A, which suggested two broad subsets despite the absence of a single overall relationship between ZGA timing in hpf and in cell cycles across species. Mammals, two nematode species and one jawless vertebrate were classified as slow developers, consistent with their relatively slow early cleavage dynamics, while all remaining species were classified as fast developers. Because developmental timing lies along a continuum, these groups were treated as practical, data-driven categories rather than strict biological classes.

### Estimation of phylogenetic signal in ZGA timing

The presence of a phylogenetic signal in the ZGA timing, measured in cell cycles and analyzed as a continuous trait, was assessed on the species tree using Blomberg’s *K* and Pagel’s λ with the *phylosig* function in the R package *phytools* v.1.9-16 (Revell, 2024). Blomberg’s K ranges from 0 to infinity, with values near 1 matching Brownian-motion expectations, values below 1 indicating weaker phylogenetic signal, and values above 1 indicating stronger phylogenetic signal. Pagel’s λ typically ranges from 0 to 1, with 0 indicating phylogenetic independence and 1 indicating Brownian-like phylogenetic covariance. The significance of Blomberg’s *K* was evaluated by 10,000 randomizations of trait values across the tips. Pagel’s λ was estimated by maximum likelihood, and its significance was assessed by a likelihood-ratio test against a model of phylogenetic independence (λ = 0).

### Inference of ancestral ZGA timing

Ancestral reconstruction of ZGA timing was performed on the time-calibrated phylogeny, treating ZGA timing as a continuous trait. Continuous ancestral states were visualized with *contMap* in the R package *phytools* (Revell, 2024), which reconstructs internal node values by maximum likelihood and interpolates trait values along branches under a Brownian-motion model (Fig. S2B). Internal node estimates and 95% confidence intervals were obtained with *fastAnc*, also implemented in *phytools* (Revell, 2024), under the same model.

### Estimation of egg volume

Egg size was estimated by measuring egg diameter and converting this value into volume. Spherical eggs were modelled as spheres (Volume = (πd^3^)/6, where d is egg diameter), and elongated eggs were modelled as prolate spheroids (Volume = (πLW^2^) /6, where L is egg length and W is egg width). When direct measurements were unavailable in the literature, egg dimensions were obtained from published microscopy images using *ImageJ* v1.54g (Schneider et al., 2012). For species in which yolk is highly concentrated in a distinct, spatially localized region of the embryo rather than distributed throughout the egg (chicken, zebrafish and medaka), embryonic cellular volume was estimated from the non-yolk region only. See Table S4 for all estimated cellular volumes as well as associated references.

### Association between ZGA onset timings and biological features

Differences in ZGA timing among categorical biological groups were tested using phylogenetic ANOVA with *phylANOVA* in *phytools* (Revell, 2024), with significance assessed using 10,000 simulations assuming Brownian motion. Associations between ZGA timing and continuous predictors were evaluated using both ordinary least squares (OLS), to obtain R^2^ values (R^2^_OLS_), and phylogenetic generalized least squares (PGLS) models, to estimate p-values (P_PGLS_) since they account for phylogenetic non-independence. PGLS models were fitted using two independent approaches: (i) the *gls* function from R package *nlme* v3.1-155, assuming Pagel’s λ correlation structure as specified in R package *ape* v5.7-1 function *corPagel* and estimating λ from the data from an initial value of 0.5 (Paradis and Schliep, 2019), and (ii) the *pgls* function from R package *caper* v1.0.3, assuming Brownian motion correlation structure. For completeness, for the nlme::gls PGLS models, we also report in Supplementary Dataset 4 the model fits using the rr2 package (v1.1.1), which provides pseudo-R2 measures. Specifically, we calculated R2_resid, R2_pred, and R2_lik from the fitted gls models using rr2::R2(). We also report the adjusted R^2^ from caper::summary.pgls as a pseudo-R^2^ for PGLS, because it quantifies variance explained relative to a null model under the same fixed phylogenetic covariance structure. Associations were additionally evaluated using phylogenetically independent contrasts with the function *pic* from the R package *ape* v5.7-1 (Paradis and Schliep, 2019). For each analysis, we provide regression coefficients, test statistics and p-values in Supplementary Dataset 4.

### Computation of ZGA-related scores

From each species’ gene expression matrices across developmental stages (see Supplementary Dataset 1), we first selected all genes with a minimum expression of 1 TPM and a maximum expression of 5,000 TPMs across stages. We then renormalized the TPMs of the remaining genes to sum 1,000,000. For each species, we computed three scores per gene, one reflecting expression changes at ZGA and two additional ones summarizing post-ZGA dynamics. All scores are available in the Supplementary Dataset 5.

i. The **ZGA score** quantifies the magnitude of transcriptional activation at ZGA, capturing both sharp onsets and more gradual activations. Sharp onsets were identified when MAT and ZGA_pre_ expression was low and expression at ZGA was high, using two tiers of fixed thresholds: either (ZGA_pre_ < 1 TPM or MAT < 1 TPM) together with ZGA ≥ 10 TPM, or a more relaxed combination (ZGA_pre_ < 5 TPM or MAT < 5 TPM) together with ZGA ≥ 25 TPM. For sharp onsets, the ZGA score was computed as:

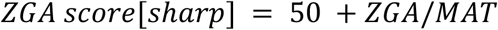

For other cases, the score aggregated information from multiple expression comparisons around ZGA, giving greater weight to the ZGA-centered comparisons (i.e., ZGA vs MAT and ZGA_pre_) and additional weight to immediate post-ZGA behavior (i.e., ZGA_+1_ vs MAT and ZGA_pre_):

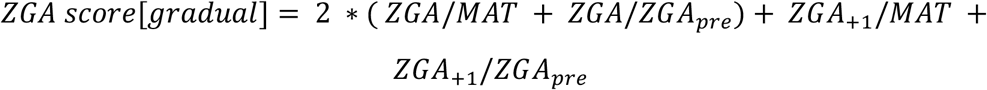
ii. The **ZGA+1 score** assesses up-regulation at ZGA_+1_ relative to ZGA, using the same rationale and two-tier thresholds:

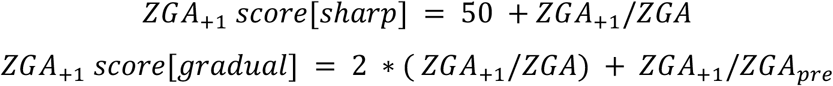
iii. The **ZGA+2 score** assesses up-regulation at ZGA_+2_ relative to ZGA_+1_, using the same rationale and two-tier thresholds:

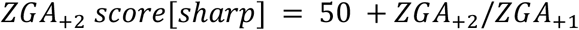

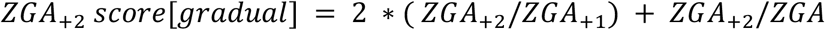

All scores were constrained to the 0–100 range and used smoothed ratios with +1 offset in numerator/denominator (not reported in the formulas above). This stabilizes ratios at low abundance and avoids division by zero. For the species for which the ZGA_+2_ stage was not available (5/61 species: *Bgr, Bbu, Oar, Csa, Cpl*) the ZGA_+2_ score was not computed.

### Definition of ZGA-related gene sets

Using these scores or the expression profiles, we defined non-overlapping gene sets in all species.

- **MAT genes**: all genes with a strong maternal contribution (MAT expression ≥ 50 re-normalized TPM).
- **MAT**_**top**_ **genes:** top 500 genes with the highest re-normalized expression at MAT stage. This gene set was used for the analysis of regulatory conservation since the same number of genes is required for a fair comparison of percentages across sets (i.e., ZGA vs MAT genes).
- **ZGA genes**: top 500 genes with the highest ZGA_score_.

### Computation of genomic features

For each species, we generated a reference GTF containing one representative transcript for each protein-coding gene, the one with the longest CDS sequence. For each gene/transcript in the reference GTF, we computed the number of introns, the total gene length (defined as the distance from gene start to gene stop in the gene entry), the 3′ UTR length (summed length of all 3′ UTR exon entries), the codon usage bias (CUB; using the SCUO method with default parameters from the R package *coRdon* v1.27.0 (Elek et al., 2025)), the GC content proportions, and the mean intergenic distance (average of the upstream and downstream distances to the closest genes). Results involving comparisons of these genomic features between MAT genes, ZGA genes, and all genes with expression ≥ 1 TPM in at least one stage are shown in Fig. 4D and Fig. S5C.

### Gene Ontology (GO) annotations

We ran *eggNOG-mapper v2*.*1*.*12* (Cantalapiedra et al., 2021) on the reference proteomes of each of the 61 species, generating uniform functional annotations across the phylogeny. In order to reduce the redundancy, we used the human GO annotation to collapse overlapping GO categories. GO categories containing more than 1,000 genes were excluded from the merging procedure and automatically included in the final annotation. We started from the smallest GO categories, systematically comparing them with increasingly bigger categories: if all genes in a smaller category also belonged to a larger one, the smaller category was removed. This left us with 2,172 human GO categories, which we then used as a template to independently reduce the GO annotation of each of the species.

### Gene Set Enrichment Analyses (GSEA)

We ran *fgsea* in R (Korotkevich et al., 2021) (with the option scoreType=“pos”) on the percentiles of genes computed on (i) decreasing expression at MAT stage or decreasing (ii) ZGA / (iii) ZGA_+1_ / (iv) ZGA_+2_ scores. These settings allow us to capture global enrichments for highly expressed MAT genes or genes increasing expression at ZGA, ZGA_+1_ and ZGA_+2_ without setting arbitrary cutoffs on the numbers of considered genes. For each species, we only tested GO categories from the GO annotations previously generated containing a minimum of 10 and a maximum of 1,000 genes. Significant GO categories in at least 60 species (i), 45 species (ii and iii) and 40 species (iv) were considered universally enriched and displayed in Fig. 4E and Fig. S6.

### Summarized expression plots

The summarized expression plots (Fig. 4A,F) were generated by first z-scoring the log_2_(TPM + 0.01) expression values across developmental stages for each gene within each species. Mean z-scores were then calculated within each gene set/category per species, and the across-species mean and standard deviation were plotted. Developmental stages were aligned across species relative to the ZGA stage, set to 0.

### Definition of genomic (G-) and regulatory (R-)conservation for ZGA and MAT genes

We first ran *Broccoli v1*.*2* (Derelle et al., 2020) on the proteomes obtained from the reference GTFs (see “Computation of genomic features”) of the 61 animal species. We obtained 28,578 gene orthogroups from which we removed lowly expressed genes (maximum expression across stages < 1 TPM). Files are available in Supplementary Dataset 7.

We then computed an estimate of the G-conservation for ZGA genes, MAT genes and the relative background (all protein-coding genes other than ZGA or MAT genes). For this purpose, we carried out bi-directional, species-pairwise comparisons starting from the previously defined gene orthogroups (see scheme in Fig. S7A). A query species gene was considered G-conserved in a target species if it was included into a gene orthogroup containing at least one ortholog from the target species. For each query-target species pair, we calculated the log2 ratio of the proportion of G-conserved genes in the ZGA or MAT gene set relative to the corresponding proportion in the background set. The full distributions of these log2 ratios for the ZGA and MAT gene sets (i.e., 60 values for each query species) are shown in Fig. S7B,C. The median of each distribution is shown in Fig. 5A.

Finally, we characterized R-conservation for ZGA and MAT_top_ genes. The MAT_top_ gene set was chosen over the MAT set for this particular analysis to avoid biases arising from different numbers of genes (500 genes in each set). A query species ZGA / MAT_top_ gene was considered R-conserved in a target species if it belonged to a gene orthogroup containing at least one target species ortholog that was itself a ZGA / MAT_top_ gene in that species. Percentages of R-conserved ZGA and MAT_top_ genes are reported in Fig. 5B,C.

### Phylostratigraphic analysis of ZGA and MAT gene sets

Gene ages were assigned with *genEra v1*.*4*.*1* (Barrera-Redondo et al., 2023). *genEra* estimates the earliest detectable evolutionary origin of protein-coding genes or gene families by genomic phylostratigraphy. We required a minimum query coverage of 50% (*--query-cover* 50) and all other settings were left as per default. Phylostratigraphic profiles of ZGA and MAT gene sets were then compared with the genomic background (see above), and enrichment was expressed as the negative log2-transformed mean phylostratigraphic rank relative to background expectation, where positive values denote enrichment in older genes and negative values denote enrichment in younger genes.

### Definition of group-specific and universally-shared ZGA and MAT gene orthogroups

In the context of these analyses, ZGA regulation indicated both ZGA genes or genes whose ZGA score fell within the top 30th percentile, while MAT regulation indicated both MAT_top_ genes or genes whose expression at the MAT stage fell in the top 30th percentile. Group-specific ZGA/MAT gene orthogroups were defined as orthogroups that (i) contained at least one ZGA/MAT_top_ gene in the queried group, (ii) showed ZGA/MAT regulation in at least 50% of the species in the queried group and (iii) did not show ZGA/MAT regulation in any other group. Universally-shared ZGA/MAT gene orthogroups were defined as orthogroups containing (i) at least one ZGA/MAT_top_ gene and (ii) at least 50% of the species in all groups with ZGA/MAT regulation. Only gene orthogroups with 250 genes or less were considered for both these analyses.

### GO enrichments for group-specific and universally-shared ZGA and MAT gene orthogroups

Since GO enrichments were performed at the orthogroup level, we generated orthogroup-based GO annotations by assigning a GO term to an orthogroup when ≥ 20% of its genes shared that annotation. GO enrichments were tested using a hypergeometric test in R *phyper(k - 1, K, N - K, n, lower*.*tail = FALSE)*, where *k* is the number of annotated orthogroups in the tested set, *K* in the background, *n* the size of the tested set, and *N* the size of the background. Backgrounds comprised all orthogroups for universally-shared ZGA/MAT analyses and all orthogroups conserved in ≥ 50% of species of the queried group for group-specific analyses. GO enrichments were considered significant for group-specific orthogroups at *p-value* ≤ 0.01 and *k* ≥ 10 (Mammalia) or *k* ≥ 5 (other groups), and for universally-shared orthogroups at *p*-value ≤ 0.01, *k* ≥ 0.2*n* (with k>1), and 50 ≤ *K* ≤ 3000. Significant enrichments are shown in Fig. S9.

## Author contributions

Conceptualization, MI; Sample acquisition, SB and HE (Bla), YL, AC-B and JMM-D (Cgi, Cte, and Ofu), SC and EH (Che), JP (Dre), Y-JL and NS (Lan), FS and JP-A (Lca), ED’A and SD’A (Mga), MIg and AS-P (Nve), JC and JRM-M (Ola), FC, MIA and SD’A (Pli), AS and APM (Pte), MIA and SD’A (Spu), Y-HS (Lpi and Pfl), RA and SD’A (Pmi), IMLS (Sro), TS-S and IA (Cdi), GQ-A (Hex), JC, XF-M and DM (Tca and Bge), KG and AB (Cpl), MH-T, CZ and RJS (additional species). Methodology, JP, CRM, IC-B, FM; Data Curation: IC-B; Software: IC-B, FM, LPI, MI; Investigation, IC-B, FM, MI; Writing, MI, IC-B and FM with feedback from the other authors; Figure generation, IC-B, FM; Funding Acquisition, MI; Supervision, MI.

## Acknowledgements

We thank members of the Irimia Lab for their critical reading of the manuscript and useful discussions. We thank Drs Alvaro Roura, Oleg Simakov, Thea Rogers and Michalis Averof for their early contributions to the study. The research has been mainly funded by the Spanish Ministry of Science and Innovation (PID2020-115040GB-I00 to MI). CRG acknowledges support of the Spanish Ministry of Science and Innovation through the Centro de Excelencia Severo Ochoa (CEX2020-001049-S, MCIN/AEI/10.13039/501100011033), and the Generalitat de Catalunya through the CERCA program. IC-B and FM were funded by FPI grants. Most in-house RNA-seq samples were generated at the CRG Genomics Unit. ChatGPT was used for minor text corrections and improvements.

## Data and Code availability

Supplementary Datasets and original code are available at Zenodo (https://doi.org/10.5281/zenodo.19227960). All raw sequencing data are available under NCBI project no. PRJNA1402383.

## Supplementary figures

**Fig. S1:**
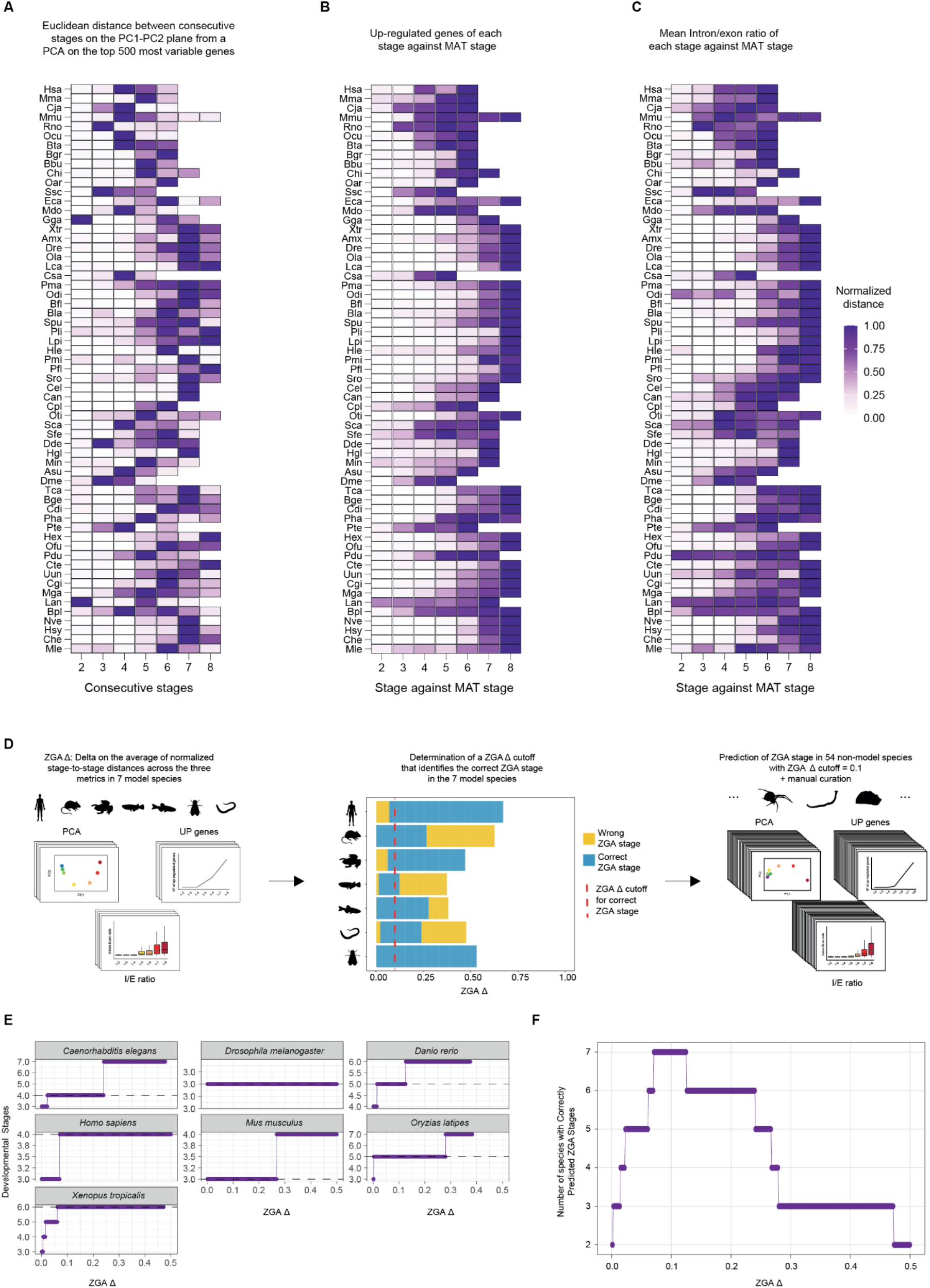
Detection and automated inference of ZGA onset across species. **A-C**. Heatmaps showing, for each species and developmental stage, the normalized value of the three transcriptomic metrics used to detect ZGA onset: **A**. Variance-weighted Euclidean distance between consecutive developmental stages in the PC1-PC2 plane from a PCA performed on the rlog-transformed expression of the top 500 most variable genes. **B**. Number of newly transcribed and strongly upregulated genes at each stage relative to the MAT stage. **C**. Mean intron/exon (I/E) read count ratio at each stage relative to the MAT stage. For each metric and species, values were scaled by their maximum, yielding normalized values between 0 and 1. **D**. Schematic of the pipeline to infer ZGA onset. (i) For each species, the three normalized metrics from panels A-C were averaged to obtain a composite score at each stage. The stage-to-stage change in this composite score (ZGA Δ) was then calculated between consecutive stages. (ii) A ZGA Δ threshold was calibrated using seven model species with well-established ZGA onset. (iii) The resulting cutoff was applied to predict ZGA stage in the remaining 54 non-model species, followed by targeted manual curation where required. **E**. Calibration plots for the seven model species, showing the predicted ZGA stage as a function of the ZGA Δ cutoff; the dashed horizontal line indicates the previously reported ZGA stage. **F**. Number of test model species for which the correct ZGA stage was recovered across ZGA Δ cutoffs. A cutoff of 0.1 yielded the highest concordance (100%) and was used for ZGA stage inference across the full dataset. Species-specific metric profiles and summary tables are provided in Tables S3.

**Fig. S2:**
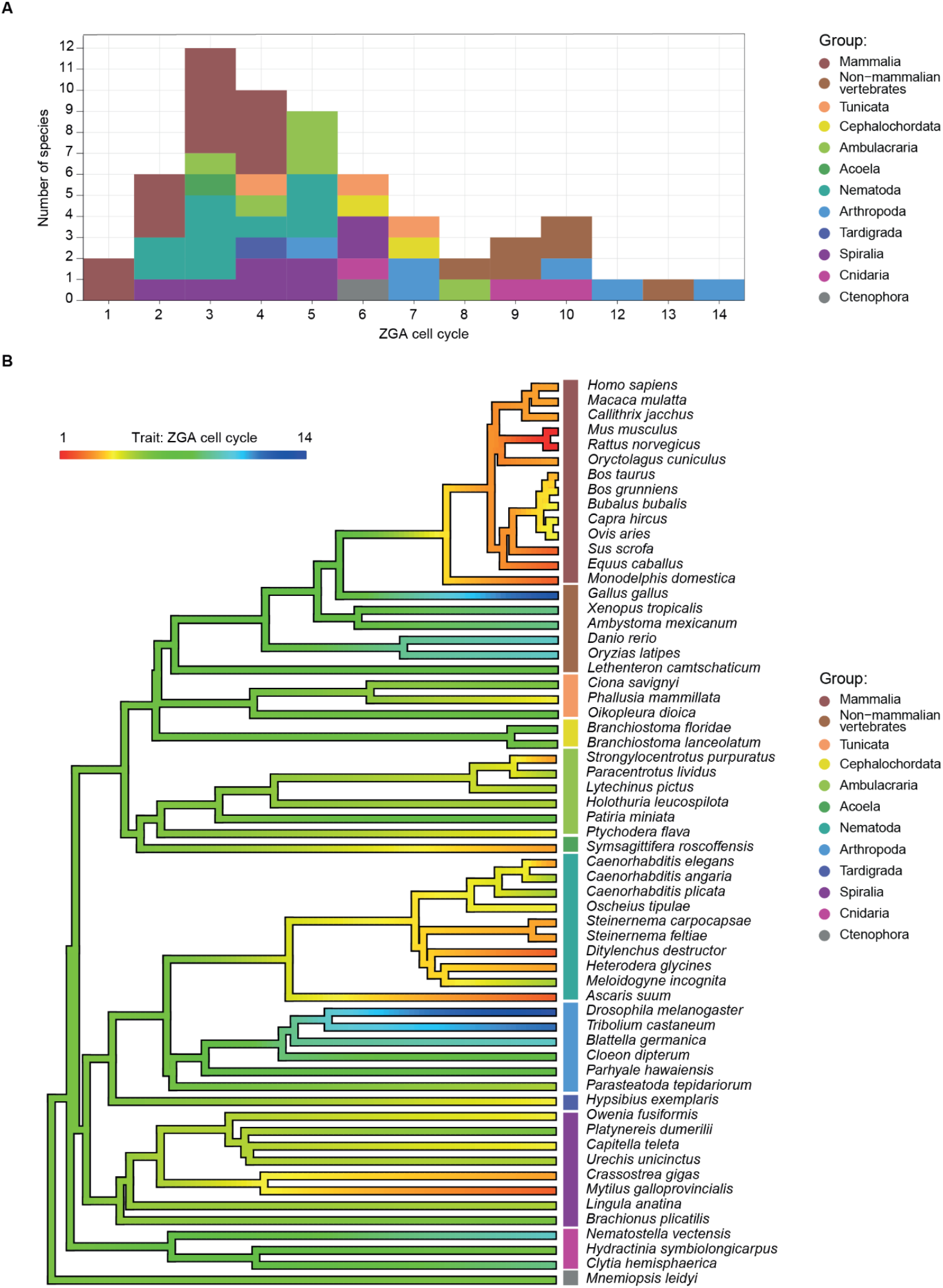
Phylogenetic distribution of ZGA onset and ancestral state reconstruction. **A**. Number of species (y axis) at each ZGA timing expressed in cell cycles (x axis), with colors indicating the relative animal group. **B**. ZGA timing mapped on the phylogeny as a continuous trait. Branch colors encode reconstructed ZGA onset expressed in cell cycles. Ancestral state reconstruction was performed using the *contMap* function in the R package *phytools* under a single-rate Brownian-motion assumption. Tips are colored according to the observed ZGA onset.

**Fig. S3:**
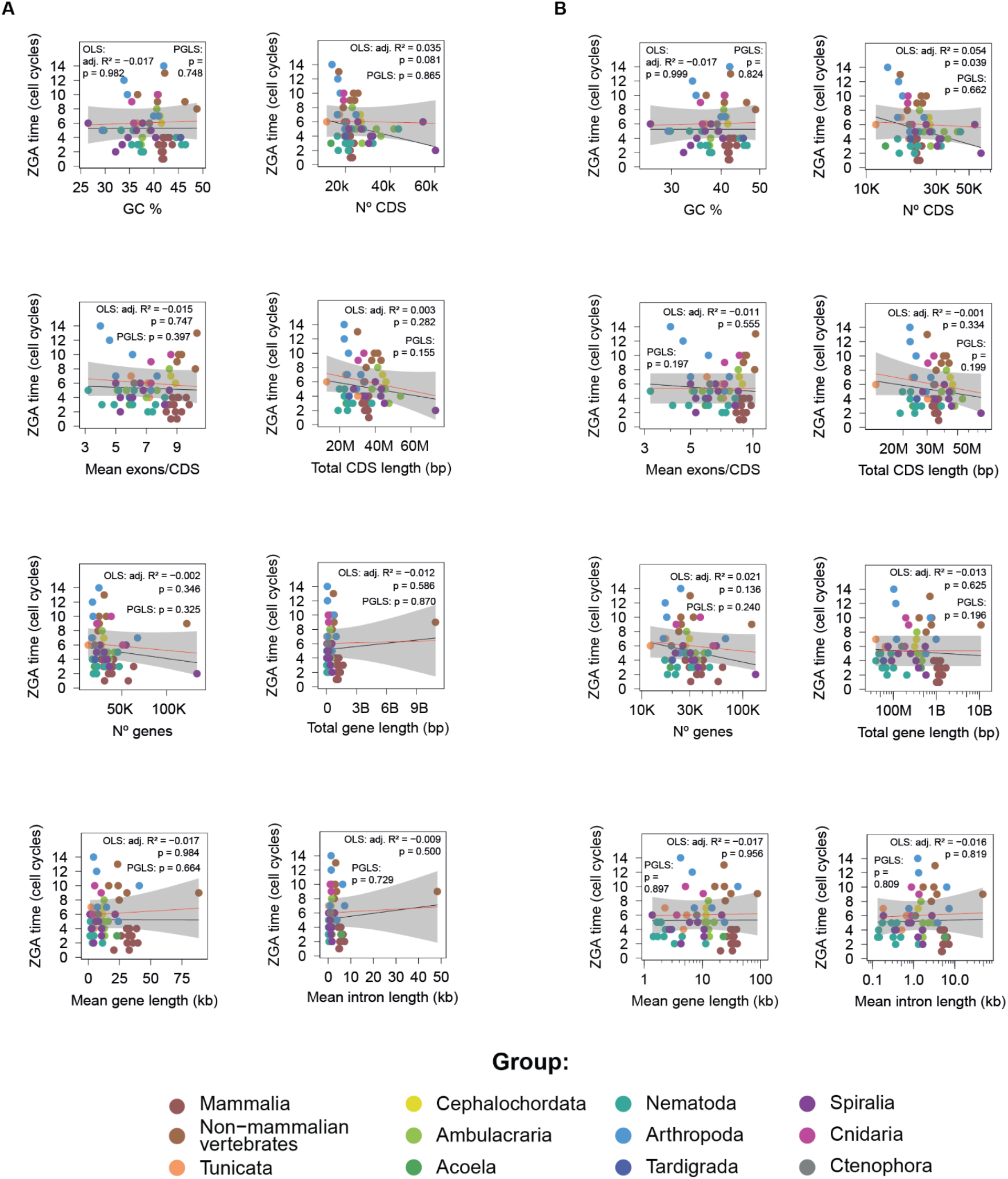
Associations between ZGA timing and continuous genomic features. **A**,**B**. In all panels, the y axis shows ZGA time expressed in cell cycles. Panel **A** shows the relationships using the original, non-log-transformed x axis values, whereas panel **B** shows the same relationships with log10-transformed x axis variables. The continuous genomic features examined are GC content (%), number of CDS, mean exons per CDS, total CDS length, number of genes, total gene length, mean gene length, and mean intron length. Black lines indicate ordinary least squares (OLS) fits, red lines indicate phylogenetic generalized least squares (PGLS) fits estimated with *nlme* and shaded ribbons show the corresponding confidence intervals for the PGLS fit. Statistics shown in each panel are the adjusted OLS R^2^ and the *P* values for both OLS and PGLS.

**Fig. S4:**
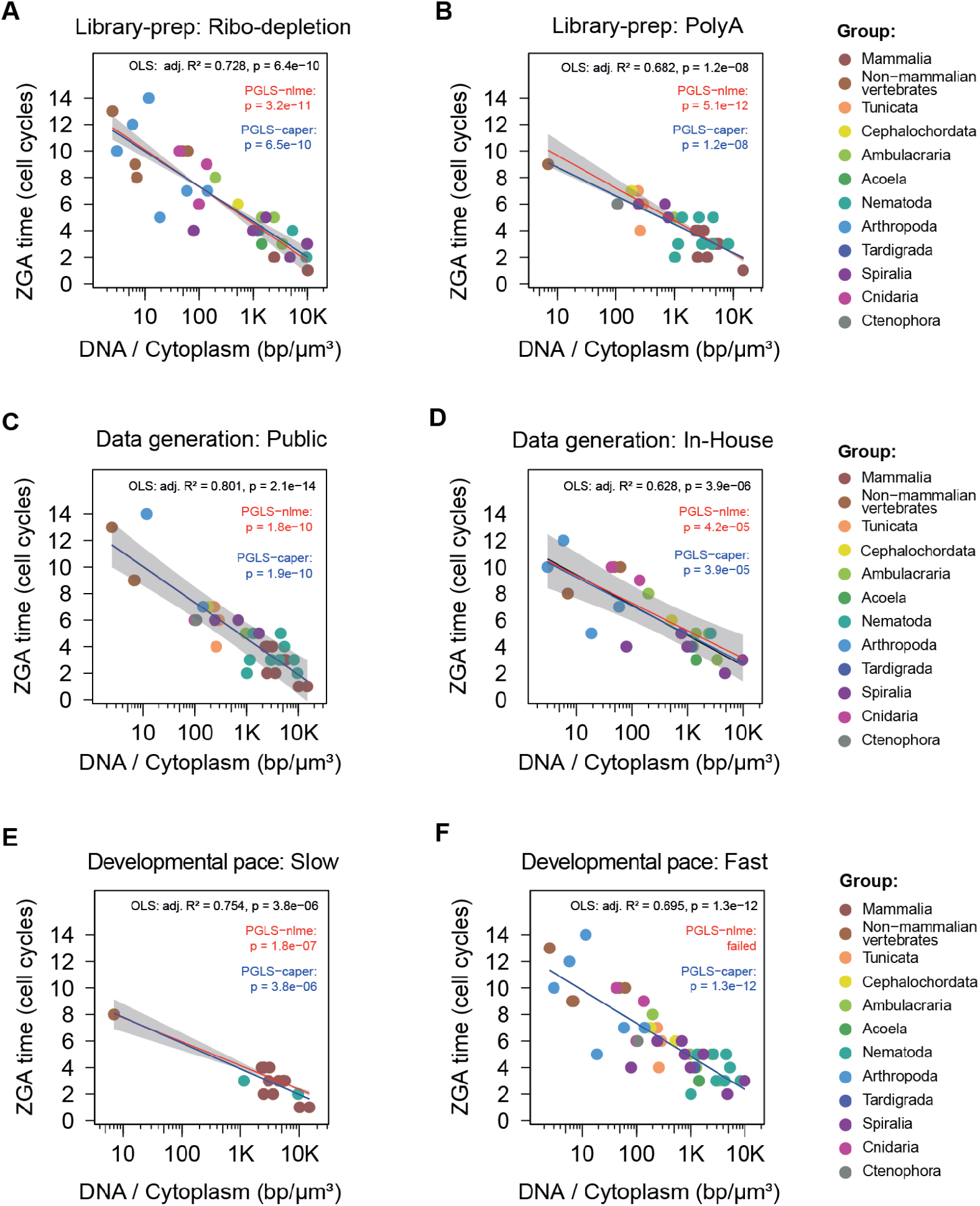
Relationship between ZGA onset and the DNA-to-cytoplasm ratio across dataset subsets. **A-F**. Plots show ZGA onset (in cell cycles) as a function of log_10_[genome size (bp) / zygote cytoplasmic volume (µm^3^)]. Species were grouped based on the library preparation method, separating ribo-depletion and PolyA selection (B); based on the data generation source, separating public data (C) and in-house generated data (D) and based on the developmental pace, separating slow developers (E) from fast developers (F). Points represent species and are colored by group. Black lines indicate ordinary least squares (OLS) regressions, red lines indicate phylogenetic generalized least squares (PGLS) fits estimated with *nlme*, and blue lines indicate PGLS fits estimated with *caper*. Gray shading shows the confidence interval of the *nlme* PGLS fit. Statistics shown in each panel include OLS R^2^ and the corresponding p-values for OLS, PGLS (*nlme*), and PGLS (*caper*).

**Fig. S5:**
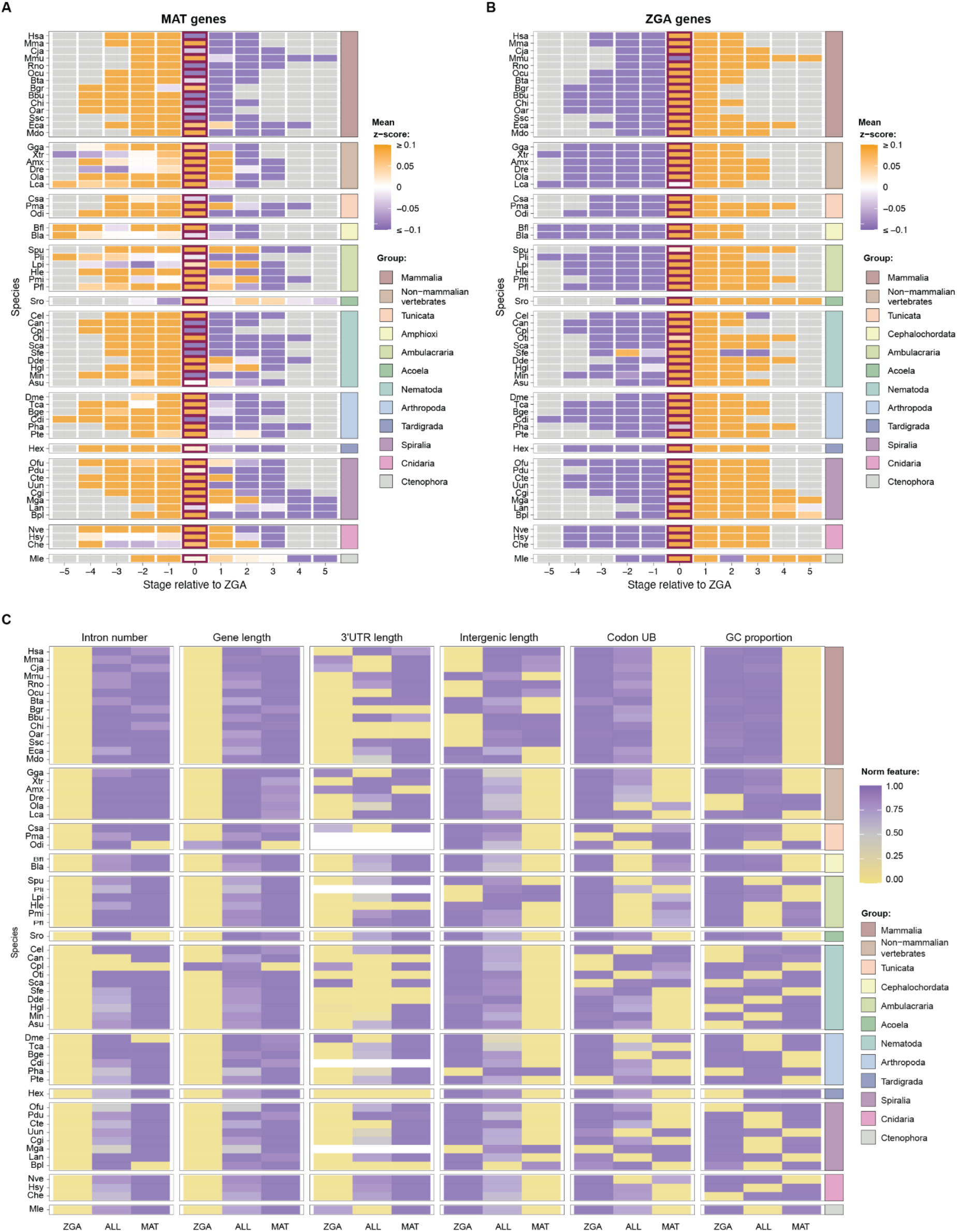
Expression profiles and genomic features across MAT and ZGA genes. **A**,**B**. Heatmaps depicting the summarized expression profiles for MAT genes (A) and ZGA genes (B), across developmental stages (x axis) and species (y axis). The log_2_(TPM + 0.01) expression values of each gene were first z-scored across stages and then averaged by species and stage. Stages were aligned relative to the ZGA stage (0). **C**. Heatmap showing within-species comparison of genomic features (x axis) between MAT genes, ZGA genes and all genes with expression TPM ≥ 1 in at least one studied stage (ALL). For each species, each feature values were normalized by the maximum observed and the lowest and highest values were set to 0 and 1, respectively. Missing values in “3′ UTR length” due to lack of UTR annotations in those species are shown in white. Abbreviations: Codon UB: codon usage bias, GC prop: GC proportion.

**Fig. S6:**
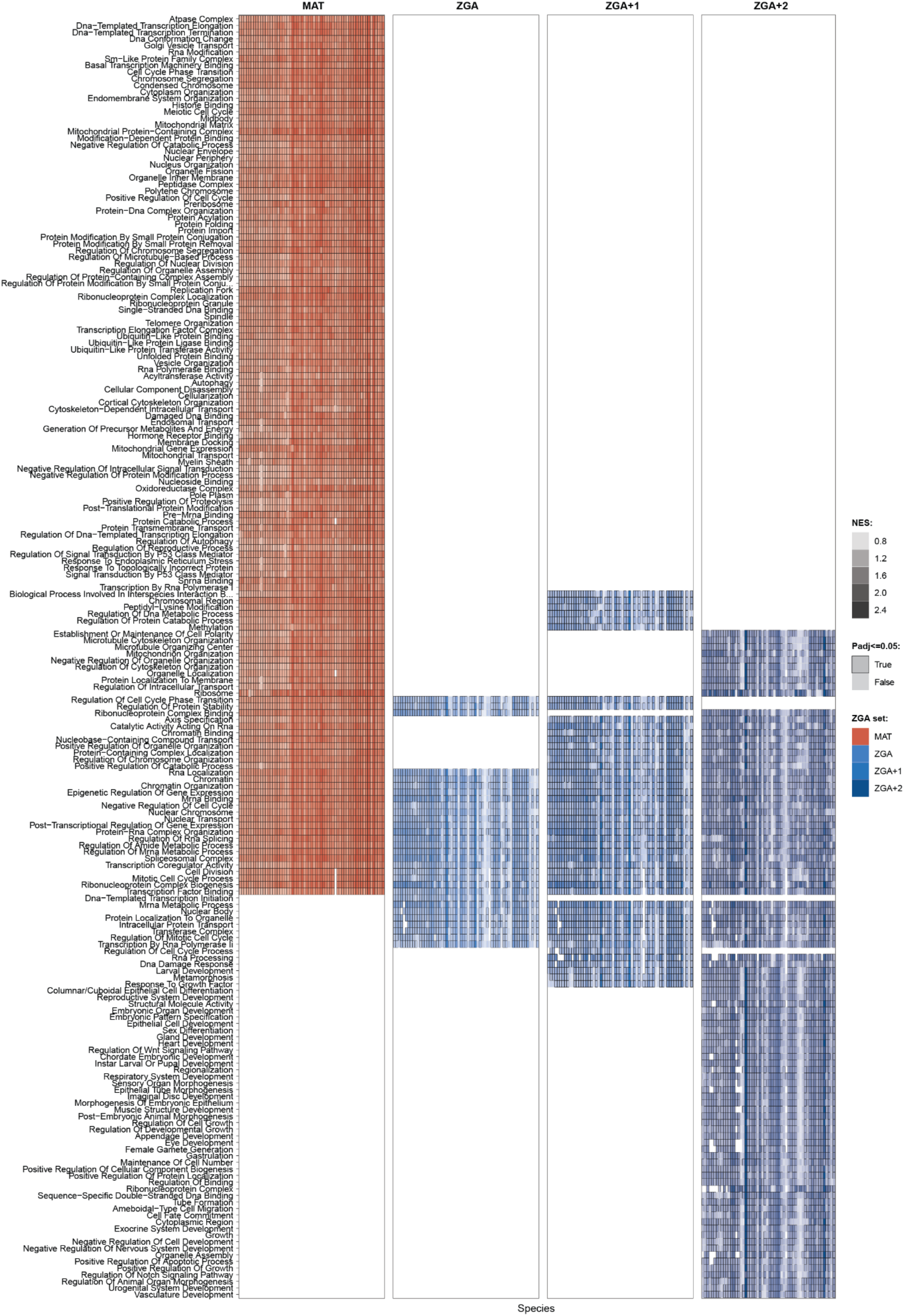
GSEA reflecting shared functions of ZGA-related sets across species. Heatmap of normalized enrichment scores (NES) for shared significant GO categories across species from GSEA performed on genes ranked by decreasing expression percentiles at the MAT stage (MAT), decreasing ZGA score percentiles (ZGA), decreasing ZGA_+1_ score percentiles (ZGA+1) and decreasing ZGA_+2_ score percentiles (ZGA+2). Only GO categories significant (adjusted *p* ≤ 0.05) in at least 60 species (MAT), 45 species (ZGA, ZGA+1), or 40 species (ZGA+2) are shown; species with significant enrichment are outlined in black. Species for which ZGA+2 scores could not be computed were excluded.

**Fig. S7:**
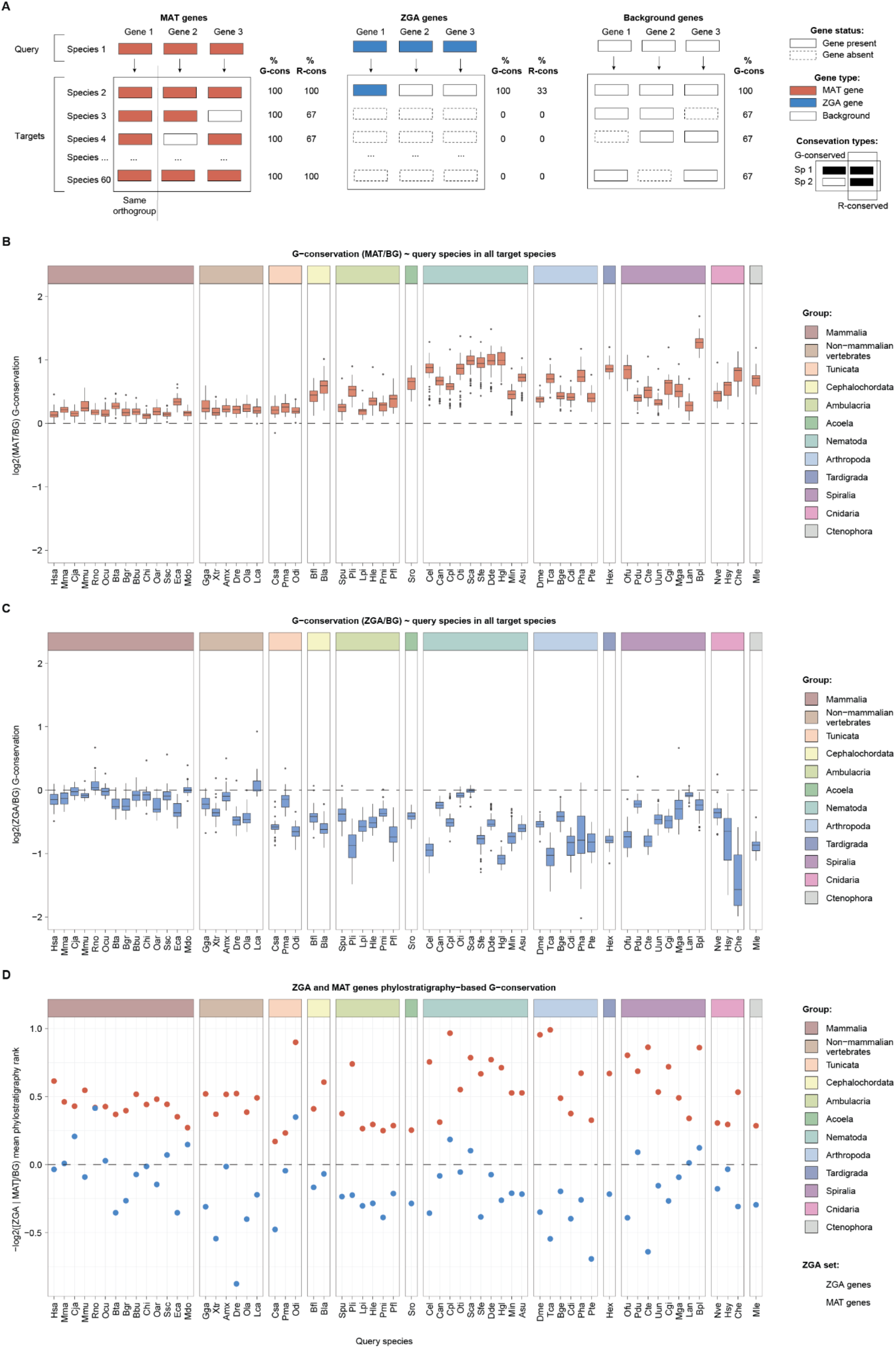
Genomic conservation of ZGA and MAT genes. **A**. Schematic representation of the definition of genomic (G-)conservation and regulatory (R-)conservation for a query species in each of the other target species. **B**,**C**. Genomic (G-)conservation (y axis) of MAT genes (B) and ZGA genes (C) in each query species (x axis) with respect to the corresponding species background. A query gene is considered G-conserved in a target species if that species contains at least one gene in the same gene orthogroup. Plotted values correspond to log_2_ ratios of G-conservation proportions between query MAT or ZGA (C) genes and the relative backgrounds in each target species. Positive / negative values on the y axis indicate more / less G-conservation than the respective backgrounds. For each query species, the background set is composed of its protein-coding genes depleted of the tested sets. **D**. Phylostratigraphic analysis of ZGA and MAT genes. Enrichment of evolutionary age for each species is plotted as the -log_2_ mean phylostratigraphic rank relative to the species background. Positive values denote an overrepresentation of ancient (conserved) genes whereas negative values denote an enrichment of younger (group-specific) genes.

**Fig. S8:**
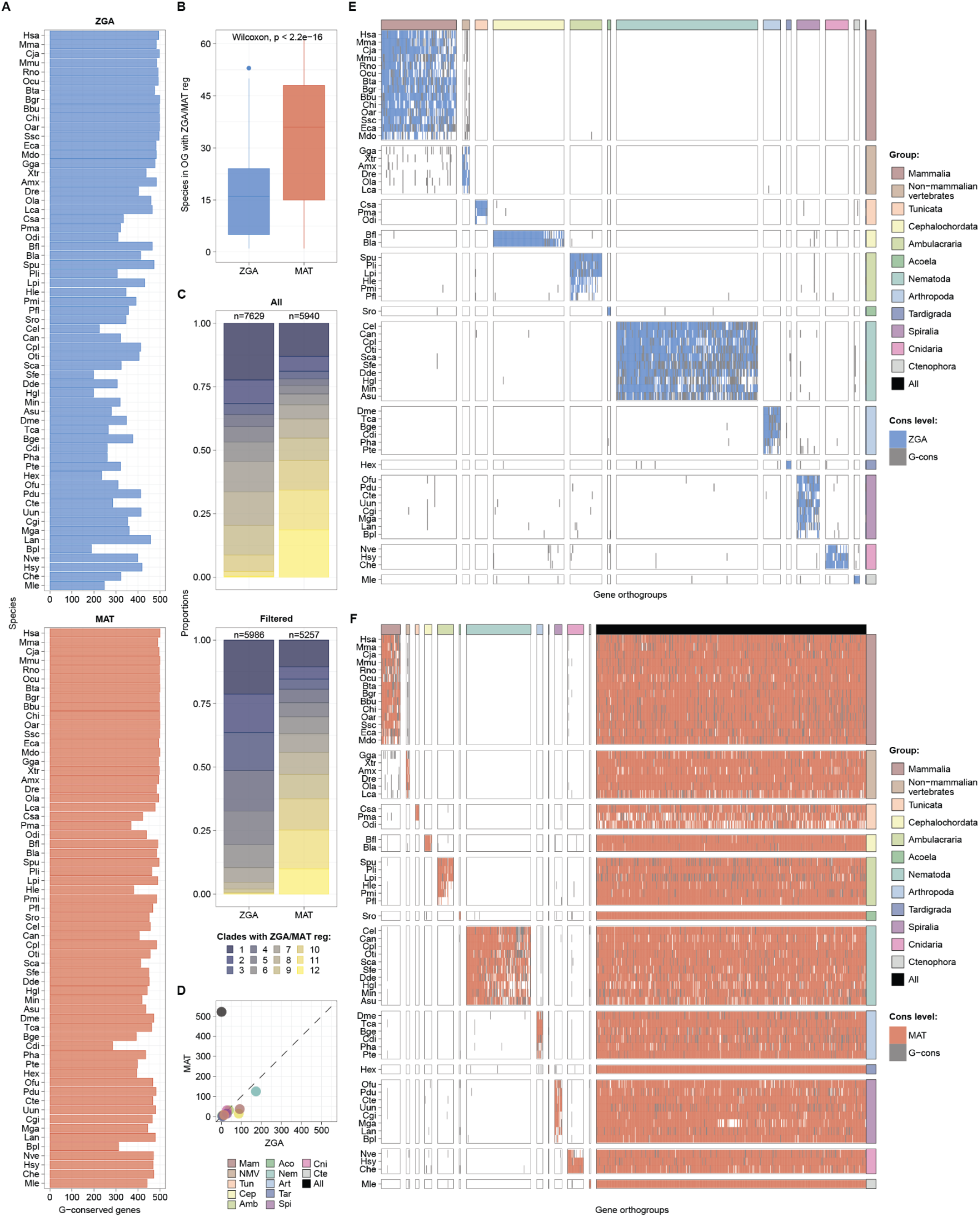
Overview of group-specific and universally-shared ZGA and MAT gene orthogroups. **A**. Number of ZGA and MAT_top_ genes (x axis) across species (y axis) that are genomically (G-) conserved in at least one other species (i.e., they have at least one detected ortholog in another species). Note that each ZGA and MAT_top_ set consists of 500 genes. **B**. Boxplots showing the numbers of species with ZGA or MAT regulation (see Methods) within gene orthogroups containing at least one ZGA gene (blue) or one MAT_top_ gene (red). **C**. Number of groups presenting ZGA or MAT regulation within gene orthogroups containing at least one ZGA or MAT_top_ gene when no filters are applied (top) or when requiring at least 50% of the species in each group to present ZGA/MAT regulation (bottom). **D**. Number of group-specific and universally-shared ZGA (x axis) or MAT (y axis) gene orthogroups. **E**,**F**. Heatmaps representing group-specific and universally-shared ZGA (E) or MAT (F) gene orthogroups (x axis) across species (y axis). Blue/red indicate ZGA/MAT regulation, whereas gray depicts those species with orthologs (G-conserved) but lacking ZGA/MAT regulation. Abbreviations: Mam: Mammalia, NMV: Non-mammalian vertebrates, Tun: Tunicata, Cep: Cephalochordata, Amb: Ambulacraria, Aco: Acoela, Nem: Nematoda, Art: Arthropoda, Tar: Tardigrada, Spi: Spiralia, Cni: Cnidaria, Cte: Ctenophora.

**Fig. S9:**
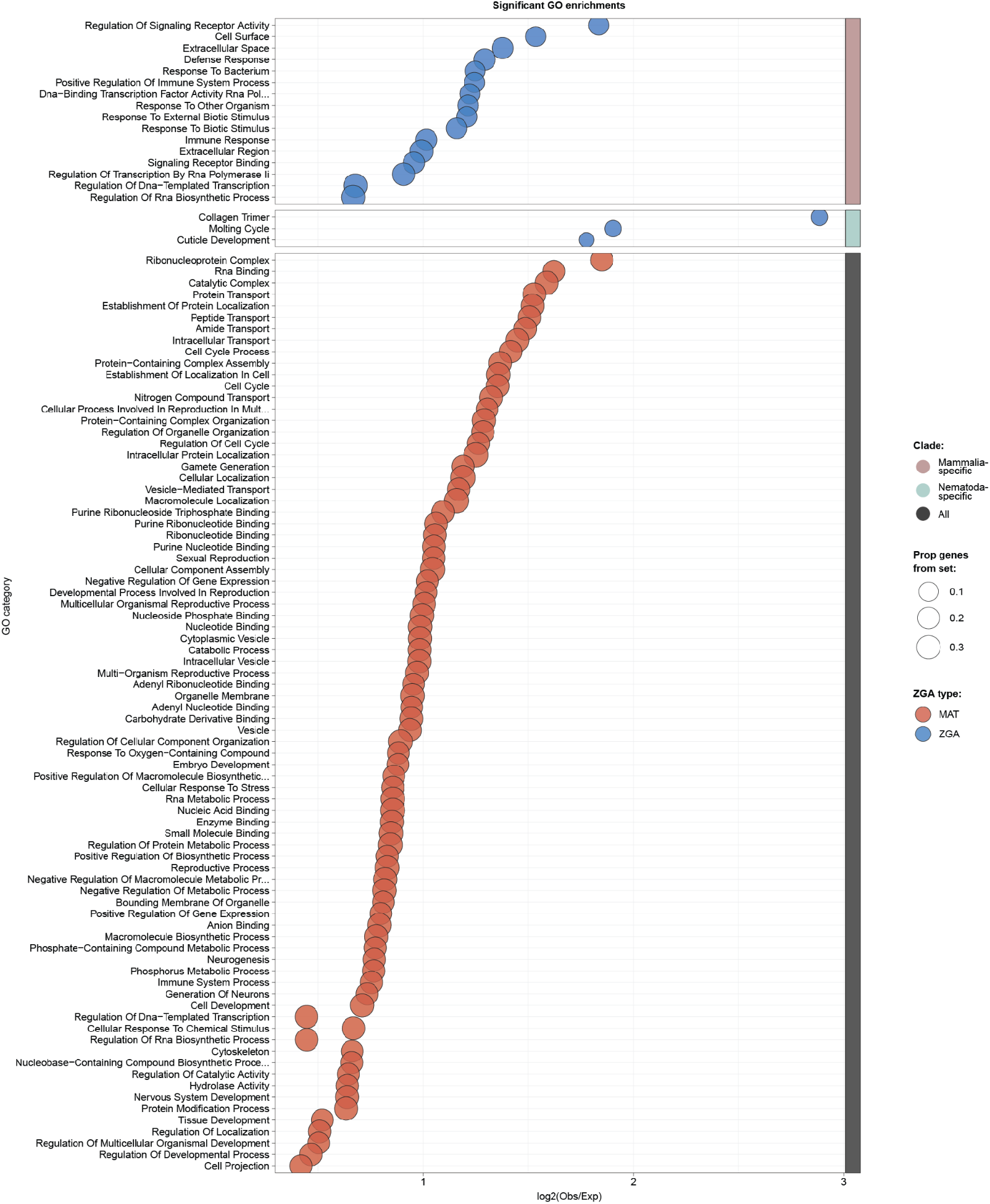
Functional enrichments of group-specific and universally-shared ZGA and MAT gene orthogroups. Significant GO enrichments of group-specific and universally-shared ZGA (blue) and MAT (red) gene orthogroups (see Methods). The size of the dots represents the proportion of gene orthogroups in each tested set annotated in the enriched GO category.

**Fig. S10:**
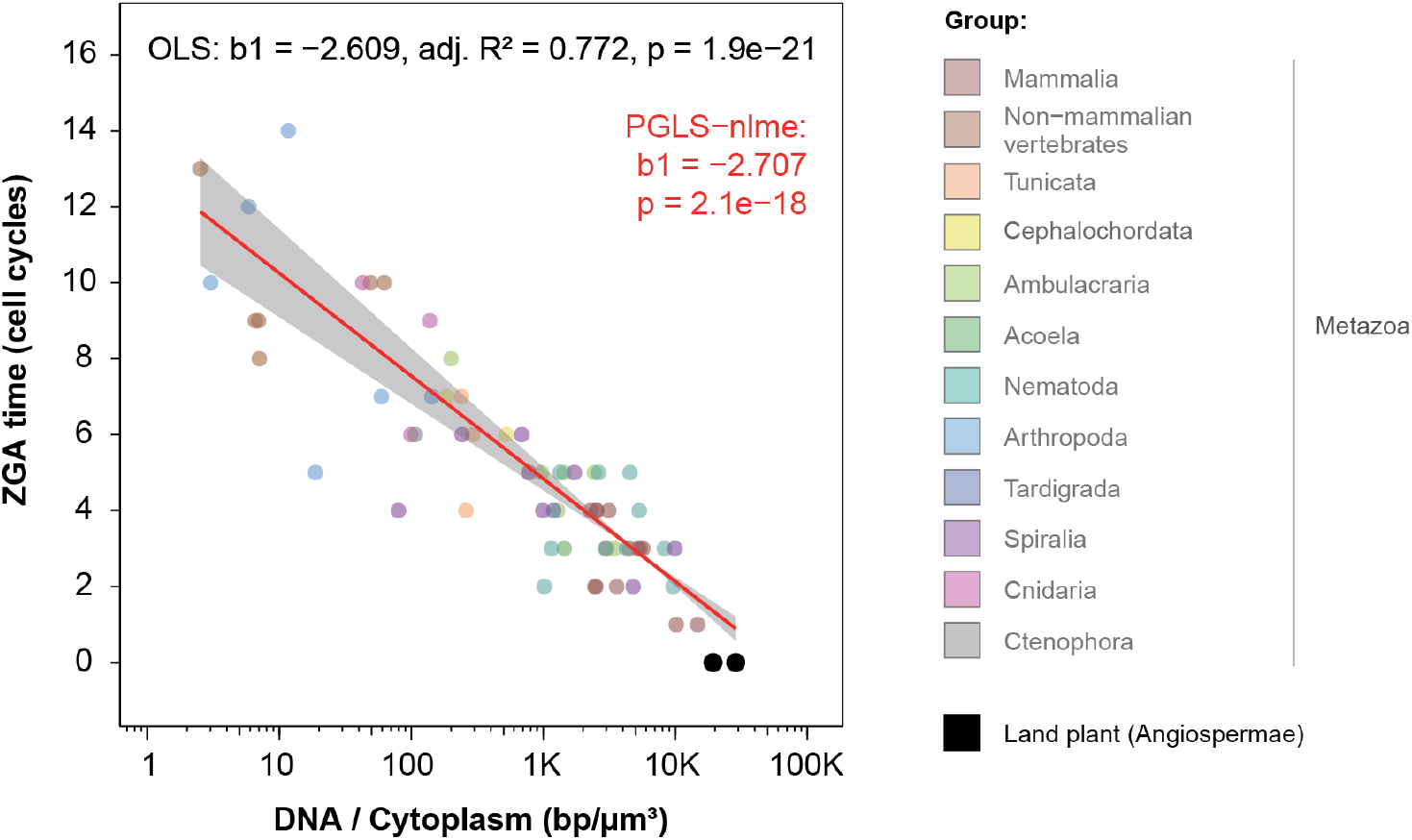
Inclusion of two angiosperms in the DNA-to-cytoplasm model for ZGA onset. ZGA timing (expressed in cell cycles) is plotted against the log_10_ DNA-to-cytoplasm ratio, calculated as the genome size (bp) divided by egg volume (µm^3^). ZGA timing for the angiosperms *Arabidopsis thaliana* and *Zea mays* was obtained from (Kao and Nodine, 2019) and (Chen et al., 2017), respectively.

## Supplementary Tables

**Table S1:** RNA-sequencing dataset metadata.

**Table S2:** Genome assembly and annotation metadata for each species.

**Table S3:** Estimated timing of ZGA onset with relative notes for the 61 animal species.

**Table S4:** Ecological, developmental and genomic traits across species.

**Table S5:** Summarized genomic features (intron number, gene length, 3′ UTR length, intergenic distances, codon usage bias and GC content proportions) for MAT, ZGA and background genes in all species.

**Table S6:** Shared GSEA enrichments across species and ZGA sets. GO categories with significant tests (adjusted *p* ≤ 0.05) in at least 60 species (MAT), 45 species (ZGA, ZGA+1), or 40 species (ZGA+2) were included.

**Table S7:** Group-specific and ancestral ZGA and MAT gene orthogroups.

## Supplementary Datasets

**Supplementary Dataset 1**: TPM expression values of protein-coding genes.

**Supplementary Dataset 2**: Exonic and intronic read counts for all samples.

**Supplementary Dataset 3**: Metrics to detect ZGA onsets and relative plots.

**Supplementary Dataset 4**: Coefficients and fit statistics of OLS and PGLS models

**Supplementary Dataset 5:** ZGA, MAT and MAT_top_ gene sets across all species, together with MAT expression, ZGA_score_, ZGA+1_score_ and ZGA+2_score_ values and relative percentiles for all protein-coding genes.

**Supplementary Dataset 6**: GSEA results across all species based on MAT expression, ZGA_score_, ZGA+1_score_ and ZGA+2_score_ percentiles.

**Supplementary Dataset 7**: Gene orthogroups across the 61 animal species.

## Supplementary Methods

In the following, we describe the protocols adopted for the generation of bulk RNA-seq samples for 23 animal species. The developmental stages sampled for each species are reported in Table S3, and the corresponding cell cycles represented in Fig. 1A.

### Lethenteron camtschaticum

Adult arctic lampreys (*L. camtschaticum*) were collected from Hokkaido, Japan. Artificial fertilization was performed in May 2021 to obtain fertilized eggs. Embryos were cultured and staged as described by (Tahara, 1988). Total RNA was extracted following (Sugahara and Pascual-Anaya, 2023), dried using GenTegra-RNA (GenTegra, Pleasanton, CA, USA), and transported for subsequent analyses. Samples collected by FS and JP-A.

### Clytia hemisphaerica

Embryos of the hydrozoan *C. hemisphaerica* were obtained by mixing gametes released from laboratory Z-strain medusae (Leclere et al., 2019). They were cultured at 18°C in 0.2 µm Millipore filtered sea water (Red Sea Salt brand). Groups of 100-200 eggs and embryos of successive developmental stages (Kraus et al., 2020) were snap frozen in lysis buffer from the Ambion RNAqueous MicroKit, kept at -80°C and later, after thawing, mRNA was purified using the same kit. Samples were then transported for subsequent analyses using GenTegra-RNA tubes (GenTegra, Pleasanton, CA, USA). Samples collected by SC and EH.

### Cloeon dipterum

Embryos of *C. dipterum* were obtained at 0, 3, 8, 13, 14.5, 23, 24, 28.5 hours after fertilisation through forced copulas (Almudi et al., 2019). Fertilised females were sacrificed and the egg sacks containing the embryos at desired stages were dissected in PBS and frozen at -80ºC in NucleoZOL (Macherey-Nagel) reagent. Total RNAs were extracted using NuceloZOL RNA extraction protocol following manufacturers instructions. RNAs concentration and quality were validated by using Qubit RNA HS Assay Kit (Invitrogen). To validate the cellular stage of the embryos, between 10 and 20 embryos approximately were separated for each dissected egg sack and fixed with 4% Formaldehyde in PBS for 1h at room temperature (RT). After 3x 10 min washes with PBTween 0.1%, embryos were stained with DAPI (1:10000) and 1:100 Alexa-Fluor 594 conjugated Phalloidin (Invitrogen) for 30 mins at room temperature., washed 3x 10 mins in PBS and mounted in glycerol 80%. Images were acquired using a Zeiss LSM 880 confocal microscope and were processed with Fiji (Schindelin et al., 2012). Samples collected by TS-S and IA.

### Symsagittifera roscoffensis

Gravid adult acoels *S. roscoffensis* were collected in Roscoff, France, in March 2024 at low tide, and transported to the laboratory, where they were maintained as per (Pennati et al., 2024). Synchronously developing embryo clutches (“cocoons”) were collected at 2 or 4-cell stages, manually triaged under the stereo-microscope, and allowed to develop in filtered sea water (FSW) in incubators in the dark at 14ºC or 18ºC until the desired stage was achieved. Clutches were dissected with fine tungsten needles and forceps and embryos stored directly in RNAlater at 4ºC overnight, then at -20ºC. A small subset of each age-matched clutch/stage was fixed in 4% PFA in PBS for 2 hours at RT, washed 3x in PBS, and stored in either PBS at 4ºC or 70% ETOH at -20ºC. The latter were used to confirm staging and estimate cell counts via immunohistochemistry/staining as follows: 3-6x 20’ washes in TBT (Tris Buffered Saline pH7.6 + 0.1% Tween); 1 hour permeabilization at RT in TBTx (1%Triton); 2 hours blocking in TBTx (0.2% Triton) +2% heat-inactivated sheep serum at RT; overnight incubation at 4ºC on the shaker in the dark with 1:200 Alexa-Fluor 568 conjugated Phalloidin (Thermo Fisher Scientific) in TBTx (0.2% Triton); 3-6x 20’ washes in TBT; 1 hour incubation on shaker at RT with 1:5000 DAPI stock (1mg/ml, Sigma) in TBS; 3-6x 20’ washes at RT in TBT. Embryos were mounted on Permafrost slides in Prolong Diamond antifade mountant (Thermo Fisher Scientific) and stored at 4 in the dark until imaging. Images were acquired using a Zeiss LSM 880 confocal microscope and were processed with Fiji (Schindelin et al., 2012). For RNA extraction, embryos were homogenized in TRIzol (Thermo, 15696018) using a MiniBeadBeater (Biospec Products) for 40 seconds prior removal of RNAlater (AM7020), along with the addition of glass beads (Merck, G8772-100G). RNA extraction was conducted using Phasemaker™ Tubes (Invitrogen, Cat. #A33248), following the manufacturer’s protocol. Final RNA concentrations were quantified using Nanodrop. Samples collected by IMLS.

### Oryzias latipes

The medaka *O. latipes* iCab wild-type strain was maintained under previously described experimental conditions (Vazquez-Marin et al., 2019). Animal experiments were carried out according to ethical regulations. Experimental protocols have been approved by the Animal Experimentation Ethics Committees at the Pablo de Olavide University and CSIC (license number 02/04/2018/041). Total RNA was isolated using the easyBLUE Total RNA Extraction Kit (Intron Biotechnology, Seongnam, Korea) according to the manufacturer’s protocol. Following extraction, RNA samples were stabilized and transported using GenTegra RNA tubes (GenTegra, Pleasanton, CA, USA). Samples collected by JC and JRM-M.

### Hypsibius exemplaris

Embryos of the tardigrade *H. exemplaris* (Z151 strain) were collected immediately after egg laying, corresponding to the parthenogenetic 1-cell stage. They were cultured at 20°C in 55 mm glass dishes filled halfway with spring water (Volvic) filtered through a 0.2µm mesh. Between 150 and 225 embryos were collected per developmental stage (Gabriel et al., 2007) and transferred to the lysis buffer from the NucleoSpin RNA Plus XS kit (Macherey-Nagel). mRNA was purified using the same kit, following the manufacturer’s protocol. Samples were then transported for downstream analysis using GenTegra-RNA tubes (GenTegra, Pleasanton, CA, USA). Samples collected by GQ-A.

### Tribolium castaneum

The *T. castaneum* enhancer-trap line pu11 (a kind gift from Y. Tomoyasu, Miami University, Oxford, OH) was maintained on an organic wheat flour diet supplemented with 5% nutritional yeast. Cultures were kept at a constant temperature of 29 °C in complete darkness. Embryos were collected following oviposition and incubated at 29 °C until the desired developmental stage. Dechorionation was performed using a diluted sodium hypochlorite (NaOCl) solution, which also effectively removed adherent flour particles from the egg surface. After thorough washing in phosphate-buffered saline (PBS), embryos were transferred to TRIzol™ Reagent (Thermo Fisher Scientific, Cat. #15596-026). RNA extraction was conducted using Phasemaker™ Tubes (Invitrogen, Cat. #A33248), following the manufacturer’s protocol. Samples collected by JC, XF-M and DM.

### Blattella germanica

Specimens of *B. germanica* were obtained from a laboratory colony maintained in complete darkness at 30 ± 1 °C and 60–70% relative humidity. Under these rearing conditions, embryogenesis lasts approximately 17 days, during which eggs develop within an ootheca carried by the female. As the ootheca is extruded from the female, the seam of the egg case initially emerges dorsally. Once extrusion is complete, the ootheca undergoes a 90° rotation, positioning the seam laterally; this point was designated as time zero of embryonic development. Because oothecae are not viable once detached from the female, the females were isolated and monitored until embryos reached the desired developmental stage. At that time, oothecae were manually removed, and embryos from the central region of the ootheca were carefully dissected, manually dechorionated, and transferred to TRIzol™ Reagent (Thermo Fisher Scientific, Cat. #15596-026). RNA extraction was performed using Phasemaker™ Tubes (Invitrogen, Cat. #A33248), according to the manufacturer’s instructions. Samples collected by JC, XF-M and DM..

### Parasteatoda tepidariorum

Embryos of *P. tepidariorum* were obtained from a laboratory culture at Oxford Brookes University. Adult spiders were maintained at 25°C. Upon cocoon production a few embryos were staged in halocarbon oil and then maintained at 25°C until the stage required. Staging was carried out as previously described (Mittmann and Wolff, 2012). Six stages were collected: 0 hours after laying and then 16-cell, 32-cell, 64-cell, 128-cell, 128+ cell (beginning of germ disc formation). A subset of embryos from each stage were imaged and RNA was extracted from the remaining embryos using QIAzol according to the manufacturer’s instructions (Qiagen). Samples collected by AS and APM.

### Ptychodera flava

*P. flava* mature adults and gametes were obtained following previously described methods (Lin et al., 2016). Eight stages were collected from two different parent pairs: 1-cell (unfertilized egg), 2-cell, 4-cell, 8-cell, 16-cell, 32-cell, 64-cell, and 100-cell stages (9 hpf). The fertilization rate was 60∼80%. Total RNA was extracted using Direct-zol RNA Microprep (Zymo) with a BioMasher II homogenizer (Nippi). Samples collected and sequenced by Y-HS.

### Lytechinus pictus

Adults of *L. pictus* were obtained from Amro Hamdoun at Scripps Institution of Oceanography, University of California, San Diego. Adult maintenance, gamete collection, and embryo cultures were based on published methods (Nesbit et al., 2019). Eight stages were collected from two different parent pairs: 2-cell, 4-cell, 8-cell, 16-cell, 32-cell, 64-cell, 100-cell, and early blastula stages. Total RNA was extracted using Direct-zol RNA Microprep (Zymo) with a BioMasher II homogenizer (Nippi). Samples collected and sequenced by Y-HS.

### Danio rerio

Zebrafish were grown in the PRBB facility in Barcelona at 28°C 14 h light/10 h dark cycle. Massive crosses were carried out in crossing tanks and the resulting eggs were mixed and distributed in batches of 100 embryos. Development of the embryos was carried out in Petri dishes with E3 medium with methylene blue and placed in an incubator at 28 ºC until the desired stage. At this point, development was arrested by placing the embryos in RNAlater (ThermoFisher AM7020) and stored at 4 ºC in order to preserve the RNA until extracted. When all the stages were obtained, every sample was extracted using the RNeasy Mini kit (Qiagen 74104) following the manufacturer’s protocol. Samples collected by JP.

### Lingula anatine

Spawning was induced via Dibutyryl cAMP and Temperature Elevation. The spawning protocol was adapted from (Tagawa et al., 1998), with modifications. Approximately 50 *L. anatina* individuals were maintained in 10 L aerated seawater tanks, with weekly water changes. Feeding was performed daily with 25 mL of Chaetoceros calcitrans (5×10^7^ cells/mL; Marine Tech, Sun Culture). One day prior to spawning induction, animals were starved overnight to reduce contamination from feces. On the day of the experiment, individuals were rinsed twice in FSW and placed individually in 90 × 15 mm dishes containing FSW. A total of 30 µL of 40 mM dibutyryl cyclic AMP (dibutyryl cAMP; Sigma D0627, MW 491.37, 19.65 mg/mL in PBS) was injected into one side of the gonad using a 0.5 mL syringe fitted with a 29G×½” needle (Terumo SS-05M2913). The dibutyryl cAMP solution was kept on ice and aliquoted to RT just prior to injection. Following injection, individuals were incubated at 29°C (approximately 4°C above ambient RT) for 2–6 hours. Spawning typically occurred 1.5–2 hours post-injection. If no gamete release was observed after 8 hours, the incubation was terminated and individuals were returned to the holding tank. During incubation, animals were checked every 30 minutes for spawning. FSW was refreshed at 30-minute intervals to remove feces and leaked body fluids; this temperature fluctuation may further stimulate spawning. The efficiency of this induction method ranged between 20–70%, depending on seasonal conditions. A minimum of 16–24 individuals per experiment is recommended to ensure adequate yield. Sperm viability was monitored post-spawning using a stereomicroscope. Sperm activity remained optimal for up to 4 hours post-spawning.

Eggs were collected using 9” glass Pasteur pipettes (Fisherbrand 12-678-20C) and transferred to 15 mL conical tubes, avoiding debris and feces. Eggs were washed twice by gentle centrifugation (∼20 seconds each) using a hand centrifuge (Hettich, Rotor No. 1014), and resuspended with a pipette. Sperm-containing seawater was collected in 50 mL conical tubes. Prior to insemination, sperm activity was confirmed under a stereomicroscope. Eggs were inseminated in 15 mL tubes for approximately 3 minutes. Excessive sperm concentration or prolonged exposure was avoided to reduce polyspermy and abnormal development. Post-fertilization, embryos were washed twice by hand centrifugation and then transferred to 150 × 25 mm dishes for culture in FSW at room temperature. Samples collected and sequenced by Y-JL and NS.

### Branchiostoma lanceolatum

Ripe adults from the *B. lanceolatum* species were collected by dredging at the Racou beach near Argelès-sur-Mer, France (latitude 42° 32′ 53′′ N, longitude 3° 03′ 27′′ E), and retrieved from the sand by sieving. Gametes were obtained by heat stimulation as previously described in (Fuentes et al., 2007). Sperm and oocytes were collected separately and fertilization was performed in vitro in Petri dishes filled with filtered sea water. Embryos were raised in the Petri dishes in the dark at constant temperatures (19°C) until the desired developmental stages. Embryos were collected at 2-cell, 4-cell, 8-cell, 16-cell, 32-cell, 64-cell, 128-cell, 256-cell, and blastula stages by centrifugation at 13,000 rpm for 30 seconds in 2mL tubes. After removing the sea water, they were frozen in liquid nitrogen and kept at −80°C prior to RNA extraction. Total RNA was extracted using the RNeasy Plus Mini Kit (QIAGEN) after disrupting and homogenizing the sample with a TissueLyser (QIAGEN, 30 Hz for 45 sec.). RNA was eluted in 30 μl of nuclease-free water. The concentration was measured with a Nanodrop and the integrity of the RNA was verified by agarose gel electrophoresis. Samples collected by SB and HE.

### Mytilus galloprovincialis

Sexually mature specimens of *M. galloprovincialis* were obtained from Irsvem Srl, a commercial shellfish farm (Bacoli, Napoli, Italy). Animals were mechanically stimulated to promote spawning by scraping the shells to remove adherent organisms and pulling the byssus. Approximately, 20–30 mussels were placed in a tank with Mediterranean FSW at 18°C and spread to easily monitor the spawning. When mussels began to spawn, each individual was washed and then transferred into a beaker containing 200 ml of Mediterranean FSW to isolate male and female gametes. Eggs were fertilized with an egg/sperm ratio 1:15 in a volume of 50 ml. The resulting zygotes were left to grow at 18°C in petri dish plates at a concentration of 250 eggs per ml until the developmental stage of interest in a 12 h light/12 h dark cycle. Total extraction from different developmental stages was carried out using the RNeasy Micro Kit (Qiagen). Specifically, centrifugation of defined embryonic stages was performed at maximum speed for 2 minutes, the pellet was quickly frozen in liquid nitrogen and stored at -80°C, to obtain approximately 7,500 embryos/replicate for each defined stage and processed following the manufacturer’s protocol. RNA quantification was performed with a NanoDrop (ND-1000 Spectrophotometer; NanoDrop Technologies, Wilmington, DE, USA). The 260/280 and 260/230 ratios of absorbance were used to assess the purity of RNA. The integrity of 28s and 18s ribosomal RNA was determined for all samples by electrophoresis first using 1% agarose gel and bioanalyzer. Samples collected by ED’A and SD’A.

### Paracentrotus lividus

Adult *P. lividus* sea urchins were collected from the Gulf of Naples (Italy) and housed in circulating seawater aquaria at Stazione Zoologica Anton Dohrn in Naples. Spawning of gravid animals was induced by vigorous shaking. Oocytes were collected in FSW, while sperm was stored without seawater (dry sperm) on ice. Oocytes were fertilized with sperm diluted 1:1000 in FSW. Fertilization success was confirmed by the visual inspection for the presence of the fertilization envelope. Only specimens with fertilization rates greater than 90% were further processed. Following fertilization, zygotes were rinsed twice with FSW to remove excess sperm and allowed to develop in FSW at 18° until the desired developmental stages. Oocytes and embryos were collected by centrifugation and pellets were snap frozen in liquid nitrogen and stored at -80°C. RNA was extracted using the Quick-RNA MicroPrep Kit (Zymo, R1050) following the manufacturer’s instructions (Jimenez-Guri et al., 2024; Paganos et al., 2023; Paganos et al., 2022). Samples collected by FC, MIA and SD’A.

### Strongylocentrotus purpuratus

Adult *S. purpuratus* sea urchins were collected from the San Diego coast by Peter Halmay, distributed by Patrick Leahy (Kerckhoff Marine Biological Laboratory, California Institute of Technology, Pasadena, CA, United States) and maintained at 14–15°C in circulating seawater aquaria at Stazione Zoologica Anton Dohrn in Naples. Spawning of gravid individuals was induced by vigorous shaking. Oocytes were collected in FSW, while dry sperm was stored on ice. Upon gamete collection, oocytes were *in vitro* fertilized by sperm, diluted 1:1000 in FSW. Fertilization was monitored through microscopic observation and confirmed by the presence of the fertilization envelope. Specimens with fertilization rates greater than 90% were further processed. Zygotes were rinsed twice with FSW, to remove excess sperm and reared at 15°C in FSW until the developmental time-points of interest. Because *S. pupuratus* is a species native to the Pacific Ocean, FSW denotes Mediterranean FSW diluted 9:1 with distilled H_2_O. Oocytes and embryos were collected by centrifugation and pellets were snap frozen in liquid nitrogen and stored at -80°C. RNA was extracted using the Quick-RNA MicroPrep Kit (Zymo, R1050) following the manufacturer’s instructions (Jimenez-Guri et al., 2024; Paganos et al., 2023; Paganos et al., 2022). Samples collected by MIA and SD’A.

### Patiria miniata

Adult *P. miniata* sea stars were provided by Patrick Leahy (Kerchoff Marine Laboratory, California Institute of Technology, USA) and maintained at 14–15°C in circulating seawater aquaria at Stazione Zoologica Anton Dohrn in Naples. Gonads were obtained by making a small dorsal incision on one arm of the animal. Testis was collected without seawater on ice. Oocytes were dissected from ovaries in FSW using fine curved forceps Oocytes were matured in 1 mM 1-methyl-adenine in FSW for 1 hour and maturation was confirmed by the visual inspection of the germinal vesicle breakdown. After fertilization, zygotes were washed twice in FSW and incubated at 15°C until the desired developmental stages. Due to the fact that *P. miniata* is a Pacific Ocean native species, FSW denotes Mediterranean FSW diluted 9:1 with distilled H_2_O. Oocytes and embryos were collected by centrifugation and pellets were snap frozen in liquid nitrogen and stored at -80°C. RNA was extracted using the Quick-RNA MicroPrep Kit (Zymo, R1050) following the manufacturer’s instructions. Samples collected by RA and SD’A.

### Owenia fusiformis

Adult *O. fusiformis* were collected near the Station Biologique de Roscoff and kept in an aerated aquarium at 15ºC. After removing them from their tubes, animals were sexed and oocytes were obtained by piercing the body wall. Oocytes were then washed in Artificial Filtered Sea Water (AFSW) by passing them through 150 and 70 µm meshes and left to incubate for an hour at 19ºC to mature. After germinal vesicle breakdown, oocytes were fertilised with a diluted sperm solution and washed twice thoroughly after 15 min to remove excess sperm. Zygotes were incubated at 19ºC in glass bowls and let develop until the desired developmental stage. Samples were collected with a mouth pipette, spun down in a 1.5 mL Eppendorf tube and pellets were snap-frozen in liquid nitrogen before storage at -80ºC. RNA was extracted using the NEB Monarch RNA extraction kit (New England Biolabs) following the manufacturer’s instructions (Liang et al., 2025). Samples collected by YL, AC-B and JMM-D.

### Capitella teleta

To collect *C. teleta* samples, mating bowls were prepared by separating gravid males and females in distinct bowls for four days. Males and females were combined in the night of the fourth day, prompting the formation of brooding tubes with fertilised oocytes the morning after. Tubes were carefully dissected, the embryos were transferred to a clean Petri dish with AFSW supplemented with 50 μg/mL penicillin and 60 μg/mL streptomycin changed daily to avoid bacterial growth, and let develop until the desired developmental stage. Embryos were spun down in a 1.5 mL Eppendorf tube and pellets snap-frozen in liquid nitrogen before storage at -80ºC. RNA was extracted using the NEB Monarch RNA extraction kit (New England Biolabs) following the manufacturer’s instructions (Liang et al., 2025). Samples collected by YL, AC-B and JMM-D.

### Crassostrea gigas

Gravid *C. gigas* were obtained from the fishmonger and opened in the lab with an oyster knife. Gonads were dissected with forceps and opened in a glass beaker, where the oocytes were let mature for an hour to improve synchronicity. After fertilisation, zygotes were washed twice with AFSW and let develop until the desired developmental stage. Samples were collected with a pipette, spun down in a 1.5 mL Eppendorf tube and pellets snap-frozen in liquid nitrogen before storage at -80ºC. RNA was extracted using the NEB Monarch RNA extraction kit (New England Biolabs) following the manufacturer’s instructions (Piovani et al., 2023). Samples collected by YL, AC-B and JMM-D.

### Caenorhabditis plicata

*C. plicata* (SB355 strain provided by the CGC) were cultured at 25 °C in modified nematode growth medium (NGM) containing 1% Agar and 0.7% Agarose. Embryos were isolated from gravid female worms through dissection in M9 drops on Rain-X-coated microscope slides to mitigate embryos sticking to glass, which were further washed through a series of M9 drops to remove bacterial contamination. Single embryos were collected in individual microfuge tubes containing 10uL of M9 solution, and promptly frozen in liquid nitrogen. RNA of single embryos was extracted using a modified established protocol (Green and Sambrook, 2020). Samples collected and sequenced by KG and AB.

### Nematostella vectensis

The *N. vectensis* culture used in this study was established from CH2 males and CH6 females (Hand and Uhlinger, 1992). Adult polyps were maintained at 18 °C in one-third filtered seawater (Nematostella medium, NM) and induced to spawn by combined temperature and light shock (Fritzenwanker and Technau, 2002). Fertilized egg packages were treated with 3% L-cysteine in NM to remove the egg jelly and subsequently cultured at 18 °C until the desired developmental stage. Eight time-points were sampled to capture the maternal-to-zygotic transition with sufficient temporal resolution, based on previous studies (Fritzenwanker and Technau, 2002; Helm et al., 2013): 2 hours post-fertilization (hpf) (zygote), 5 hpf (morula, 16-32 cell stage), 7 hpf (early blastula, cytokinesis 7), 10 hpf (blastula, cytokinesis 10), 12 hpf (blastula, cytokinesis 12), 21 hpf (early gastrula), 24 hpf (gastrula). Unfertilized eggs recently laid and de-jellied were used as time point 0. Only normally developing embryos were selected; unfertilized or un-cleaved embryos were excluded. Fertilization efficiency exceeded 90%. For each time point, approximately 400–500 embryos were pooled, thoroughly homogenized in TRIzol™ reagent (Thermo Fisher Scientific, Cat. No. 15596018), and stored at −80 °C. Total RNA was extracted and purified using the Direct-zol RNA Miniprep Kit (Zymo Research, Cat. No. R2050), followed by an additional TURBO™ DNase treatment and RNA cleanup (Thermo Fisher Scientific, Cat. No. AM1907). RNA integrity was assessed on an Agilent TapeStation system (average RIN = 8), and RNA concentrations were measured using the Qubit RNA HS Assay Kit (Thermo Fisher Scientific, Cat. No. Q32852). Samples collected by MIg and AS-P.

